# Role of 6-Phosphofructo-2-Kinase/Fructose-2,6-Bisphosphatase-3 in Maintaining Redox Homeostasis and DNA Repair in Non-Small Cell Lung Cancers Under EGFR-Targeting Therapy

**DOI:** 10.1101/2024.03.25.586703

**Authors:** Nadiia Lypova, Susan M. Dougherty, Brian F. Clem, Jing Feng, Xinmin Yin, Xiang Zhang, Xiaohong Li, Jason A. Chesney, Yoannis Imbert-Fernandez

**Author notes:** Authors to whom correspondence should be addressed: Nadiia Lypova, Yoannis Imbert-Fernandez.

## Abstract

The efficacy of FDA-approved tyrosine kinase inhibitors (TKIs) targeting EGFR is limited due to the persistence of drug-tolerant cell populations, leading to therapy resistance. Non-genetic mechanisms, such as metabolic rewiring, play a significant role in driving lung cancer cells into the drug-tolerant state, allowing them to persist under continuous drug treatment. This study aimed to investigate the impact of the glycolytic regulator 6-Phosphofructo-2-kinase/fructose-2,6-bisphosphatase (PFKFB3) on the metabolic adaptivity of lung cancer cells to EGFR TKI therapies. Using two EGFR-driven non-small cell lung cancer cell lines, PC9 and HCC827, we analyzed metabolic changes in cells exposed to EGFR inhibitors and evaluated the effect of PFKFB3 inhibition on metabolic adaptations during TKI treatment. Our results indicate that PFKFB3-mediated glycolysis sustains ATP production upon TKI treatment. Metabolomics studies revealed that PFKFB3 inhibition in TKI-treated cells limits glucose utilization in glycolysis, TCA cycle, and polyol pathway. Additionally, we show that pharmacological inhibition of PFKFB3 overcomes TKI-driven redox capacity by diminishing the expression of glutathione peroxidase 4 (GPX4), which in turn, exacerbates oxidative stress. Our study also revealed that PFKFB3 contributes to DNA oxidation and damage by controlling the expression of DNA-glycosylases involved in base excision repair. In TKI-treated cells, PFKFB3 inhibition reduced ATM expression and limited DNA damage repair, increasing sensitivity to DNA integrity insults.

In summary, our results suggest that inhibiting PFKFB3 can be an effective strategy to eradicate cancer cells surviving under EGFR-TKI therapy before they enter the drug-resistant state.

**STATEMENT OF IMPLICATION:** Targeting PFKFB3 can improve the efficacy of EGFR-targeting TKIs by restricting non-genetic adaptations embraced by drug-tolerant cells.

## INTRODUCTION

Non-small cell lung cancer (NSCLC) development and aggressiveness are largely driven by deregulated expression and activating mutations of EGFR (mutEGFR, 15–30% of NSCLC cases) (1). While three generations of EGFR tyrosine kinase inhibitors (TKIs) have improved progression-free survival in patients with EGFR-driven NSCLCs, they have failed to provide overall survival benefits due to tumor reoccurrence. Beyond genomics, accumulating evidence suggests that mutEGFR NSCLC cells overcome EGFR TKI-mediated cytotoxicity through parallel non-genomic resistance mechanisms arising from transcriptomic, metabolomic, and epigenetic changes (2,3). One such mechanism is metabolic remodeling, which rewires cancer cell metabolism to meet the altered bioenergetic, biosynthetic, and redox demands imposed by selective treatment pressure. The heightened metabolic dependency of EGFR-driven cancers on glycolysis underscores glucose as a key substrate involved in metabolic rewiring (4,5).

The initial response to TKI therapy triggers the excessive accumulation of reactive oxygen species (ROS), resulting in oxidative stress and a disruption in redox homeostasis. This oxidative environment promotes the oxidation of macromolecules, including DNA, leading to consequential DNA damage. Glycolysis plays a crucial role in sustaining rapid DNA repair and alleviating oxidative stress by regulating the pentose phosphate pathway in lung cancer (6,7). Metabolic rewiring, coupled with the action of antioxidant defense enzymes such as glutathione peroxidase 4 (GPX4), enables cancer cells tolerate treatment by improving their redox capacity and developing adaptive mechanisms to counteract oxidative stress.

It has been shown that the glycolytic enzyme 6-Phosphofructo-2-kinase/fructose-2,6-bisphosphatase (PFKFB3) plays a crucial role in maintaining redox homeostasis (8) and regulating DNA damage repair (6,9). These findings led us to hypothesize that PFKFB3-driven metabolism limits the efficacy of EGFR-targeting therapies by alleviating oxidative stress and preserving DNA integrity. PFKFB3 controls glycolytic flux by catalyzing the synthesis and degradation of fructose-2,6-bisphosphate (F2,6BP) (10). F26BP is an allosteric activator of 6-phosphofructo-1-kinase (PFK-1), a rate-limiting enzyme and essential control point in the glycolytic pathway (11). Conversely, decreased expression of PFKFB3 redirects glucose metabolism towards the pentose phosphate pathway, ensuring the synthesis of reducing agents such as NADPH and GSH required for cellular survival, albeit at the expense of energy management (12).

However, the precise role of PFKFB3 in maintaining redox homeostasis in cells subjected to TKI therapy remains unknown.

In previous studies, we demonstrated that constitutively active EGFR drives glycolysis, while targeted inhibition of PFKFB3 abrogates EGFR-mediated glycolysis, resulting in reduced cell viability in lung cancer cells (4). The present study aimed to investigate the mechanisms by which PFKFB3 facilitates cell survival in response to EGFR inhibitor therapies. Two EGFR-driven NSCLC cell lines were used to elucidate the role of PFKFB3 in the metabolic perturbations induced by EGFR inhibitors. Notably, pharmacological inhibition of PFKFB3 induced oxidative stress, effectively overcoming the GPX4-dependent redox capacity of TKI-treated cells and improving TKI cytotoxicity. Additionally, we found that PFKFB3 controls DNA oxidation by regulating the expression of enzymes involved in base excision repair (BER). Simultaneously, our data revealed that TKI-treated cells exhibit limited DNA damage repair, indicating DNA integrity as molecular vulnerability of the cells during therapy. Consequently, PFKFB3 inhibition in TKI-treated cells triggered oxidative stress and DNA damage, leading to ROS-dependent cell death. Our results indicate that cells undergoing EGFR-TKI therapy rely on PFKFB3 to maintain redox homeostasis and preserve DNA integrity, highlighting the potential of inhibiting PFKFB3 as an effective strategy for eradicating cancer cells tolerant to EGFR-TKI therapy.

## MATERIALS AND METHODS

### Reagents

Erlotinib (Cat# S7786) and Osimertinib (Cat# S7297) were obtained from Selleckchem. PFK158 (Cat# HY-12203) and KAN0438757 (Cat# HY-112808) were purchased from MedChem Express. Antioxidant N-acetylcysteine (#A7250) was received from Sigma. TTP was purchased from Sigma-Aldrich(#T0251); GTP (#16060), ATP (#14498), CTP (#18147), and UTP (#9003530) were obtained from Cayman Chemical.

### Cell culture

HCC827 (RRID:CVCL_2063) cells were purchased from the American Type Culture Collection (ATCC), and PC9 (RRID:CVCL_XA19) were ordered from Sigma-Aldrich. Cells were cultured in RPMI (Sigma Cat#R8758) supplemented with 10% fetal bovine serum (FBS, Clontech) and 50 μg/ml gentamicin (Life Technologies). Cells were incubated at 37°C with 5% CO2. Cell lines were authenticated prior to the experiments and cultured for no longer than 20 passages.

### Antibodies and Western blotting

Whole-cell lysates were processed using RIPA buffer (Thermo Fisher) supplemented with protease inhibitors. Protein concentration was determined using the BCA protein assay kit (ThermoFisher, #A55864) following the manufacturer’s instructions. Proteins were separated on 10% or 4-20% CRITERION TGX gels under reducing conditions and transferred to Immun-Blot PVDF membranes (Bio-Rad). The membranes were blocked with 5% BSA or 5% nonfat milk in TBS-T (0.1% Tween20) and immunoblotted with the indicated antibodies. HRP-conjugated goat anti-rabbit or anti-mouse IgG were used as secondary antibodies. Amersham ECL Prime Western blotting detection reagent (GE Healthcare) was used to detect immunoreactive bands. The membranes were visualized on autoradiography film BX (MidSci). Quantitative densitometry was performed with ImageJ (NIH, RRID:SCR_003070) using the Gel Analysis method (http://rsb.info.nih.gov/ij/docs/menus/analyze.html#gels.) The complete list of antibodies is provided in the supplemental file.

### Glycolysis assay

PC9 or HCC827 cells growing in 6-well plates were incubated in 500 µl of medium containing 1 µCi of 5- [^3^H] glucose for 60 min in 5% CO_2_ at 37ºC. The glycolysis assay was performed as described in ref. (13). Protein concentration was determined using the BCA assay according to the manufacturer’s instructions and measured on a Powerwave XS plate reader (Biotek). Counts were normalized to protein concentration. Data are presented as mean±S.E of three independent experiments with technical duplicates.

### Glucose Uptake

PC9 or HCC827 cells were treated with vehicle control, erlotinib, or/and PFK-158 for 24 hours, and glucose uptake was assessed as described before (13). Counts were normalized to protein concentration. Data are presented as mean±S.E. of three independent experiments with biological duplicates.

### [U-^13^C]-glucose tracer studies

PC9 cells were seeded in 10 cm dishes (Corning, #430167) at density 1 × 10^6^ cells and exposed to vehicle control, erlotinib, or/and PFK-158 treatments for 12 hours. Subsequently, cells were labeled for 24 hours with RPMI medium (Gibco, #11879020) supplemented with 1 g/L [U-^13^C]-glucose (Cambridge Isotope Laboratories, #110187-42-3), 10% dialyzed fetal bovine serum (R&D Systems, #S12810H), and the appropriate treatments were maintained. The cells were then washed three times in ice-cold PBS and quenched with acetonitrile. Metabolites were extracted in acetonitrile:water (1 mL:667 µL). After three freeze-thaw cycles, samples were centrifuged at 3000×g for 20 min at 4°C to separate cell debris. The supernatant and pellet fractions were separated and vacuum-dried by lyophilization. The dried supernatant fractions were collected for 2D-LC-MS/MS analysis.

### LC-MS analysis and data processing

Polar metabolites were detected using the method described previously (14). All samples were analyzed on a Thermo Q Exactive HF Hybrid Quadrupole-Orbitrap Mass Spectrometer coupled with a Thermo DIONEX UltiMate 3000 HPLC system (Thermo Fisher Scientific, Waltham, MA, USA). The LC system was equipped with a reversed phase column (RPC, Waters Acquity UPLC HSS T3 column, 2.1 × 150 mm, 1.8 µm) and hydrophilic interaction chromatography column (HILIC, a Millipore SeQuant ZIC-cHILIC column, 2.1 × 150 mm, 3 µm) configured in parallel. Each column was connected to a 2-μL sample loop, and the column temperature was set to 40°C. All samples were analyzed in a random order in positive (+) and negative (-) modes to obtain complete MS data for metabolite quantification. For the metabolite identification, unlabeled samples were analyzed by 2D-LC-MS/MS in positive and negative modes at three collision energies, 20, 40, and 60 eV. Total metabolite levels were normalized to the pellet fraction weights. Data are presented as mean±S.D. of two independent experiments with 6 biological replicates.

### Fractionation of Soluble and Chromatin-Bound Proteins

PC9 and HCC827 cells (1×10^6^) were seeded on 100 mm plates and treated the following day for 24 hours. Soluble and chromatin-bound proteins were fractionated following the protocol described in (15). Equal amounts of cells were processed for fractionation. Ten microliters of each soluble or chromatin-bound nuclear fraction were loaded on 4-20% CRITERION TGX SDS-PAGE gels (Bio-Rad, #4561085).

### PFKFB3 siRNA transfection

PC9 cells were seeded in six-well plates at a density of 12x10^4^ cells/well in 2 ml of complete medium 24 hours before transfection. Transfections were performed using Lipofectamine RNAiMAX (Thermo Fisher Scientific, #13778030) following the manufacturer’s protocol. The following siRNAs were used: control siRNA that have no homology to any sequence in the human genome were used as the controls (Thermo Fisher Scientific, #4390846); PFKFB3 siRNAs (Thermo Fisher Scientific, siRNA#1 - #HSS103358, siRNA#2 - #HSS103359). The next day, the media was changed, and cells were exposed to 0.5 µM erlotinib for 24 hours.

### ATP assay

PC9 and HCC827 cells were exposed to the appropriate treatment for 48 hours. Cells were collected and digested in 100 µl of Passive Lysis buffer (Promega, #E1941). The intracellular concentrations of ATP in the cultured cells were assayed using the ATP Determination Kit (Molecular Probes, Invitrogen, #A22066) according to the manufacturer’s protocol. ATP levels were normalized to protein concentration. The data are presented as the mean±S.E. of three independent experiments, each with technical replicates.

### GSH/GSSG measurement

PC9 and HCC827 cells were exposed to the appropriate treatment for 24 hours. Total, reduced (GSH), and oxidized (GSSG) glutathione levels in biological samples were measured using Ellman’s Reagent and glutathione reductase using Glutathione GSH/GSSG Assay (Sigma-Aldrich, #MAK440) following the manufacturer’s instructions. The data are presented as the mean±S.D. of two independent experiments, each with technical replicates.

### ROS determination

Cellular ROS levels were measured using dichlorofluorescin diacetate (DCFDA/H2DCFDA cellular ROS assay kit, Abcam, Cat# ab113851) following the manufacturer’s instructions. Briefly, live cells, exposed to different treatments, were incubated with 20 μM DCFDA for 45 min at 37°C. Cellular ROS levels were measured based on DCF fluorescence upon DCFDA oxidation by ROS. Images were taken at 4× magnification with the EVOS FL Imaging System (Thermo Fisher Scientific). Cells were analyzed using the adopted counting and scoring CellProfiler pipeline (RRID:SCR_007358). The number of cells with DCF signal above the threshold was normalized to the total number of cells per field and presented as % DCF positive cells. At least three independent fields were analyzed per treatment condition replicate. The data are presented as the mean±S.E. of three independent experiments, each with technical replicates.

### Neutral Comet assay

DNA damage was evaluated using the Comet Assay kit (Bio-Techne #4250-050-K) following the manufacturer’s instructions. Briefly, after PC9 or HCC827 cells were exposed to different inhibitors for 24 hours, 4x10^3^ cells were used to perform the Neutral Comet assay. Images of individual comets were acquired at 20 x magnification with EVOS FL Imaging System (Thermo Fisher Scientific). A minimum of 50 Comets per treatment replicate were quantified using ImageJ COMET Open software (cometbio.org). Olive tail moment was used to evaluate DNA damage. The tail moment, expressed in arbitrary units, was calculated by multiplying the percent of DNA (fluorescence) in the tail by the length of the tail in μm. The data are presented as violin plots for individual comet tails from two independent experiments, each with technical duplicates.

### Immunofluorescence Microscopy

PC9 and HCC827 cells cultured on coverslips were exposed to the appropriate treatments for 24 hours. Cells were fixed with 4% paraformaldehyde (Thermo Fisher Scientific #28908), permeabilized with 0.1% saponin (Sigma, #SAE0073), and blocked in 5% heat-inactivated goat serum (Thermo Fisher Scientific #31873) for 1 hour. After incubation with anti-γH2AX antibody (Millipore Cat# 05-636, RRID:AB_309864, 1:500) for 1 hour, cells were washed 3 times in PBS before addition of secondary antibody (Molecular Probes Cat# A-11008 (also A11008), RRID:AB_143165, 1:200) for 1 hour. Following six washes with PBS, coverslips were mounted using ProLong Gold antifade reagent with DAPI (Invitrogen, #P36931). Images were acquired with Nikon A1R confocal microscope using a 40× magnification lens with appropriate laser channels and processed in the CellProfiler pipeline (RRID:SCR_007358). At least three random fields were analyzed for each condition. Results of three independent experiments with technical replicates are presented.

8-OXO-G levels were evaluated using the 8-oxoG antibody (R and D Systems Cat# 4354-MC-050, RRID:AB_1857195). Briefly, 35,000 cells growing on coverslips were exposed to inhibitors. After 24 hours, cells were fixed and processed as described (16). Images from at least three random fields were acquired at 20 x magnification with the EVOS FL Imaging System. Cells were analyzed using the adopted counting and scoring CellProfiler pipeline. Results of two independent experiments with technical replicates are presented.

### *In silico* ingenuity network analysis

Gene expression profile, deposited by Lantermann et al. (17), was downloaded from NCBI’s Gene Expression Omnibus (GEO) database (RRID:SCR_005012) under accession number GSE67051 (accessed on January 20, 2021). The list of differentially expressed genes (DEG, RNA-seq) in the erlotinib vs. DMSO comparison for each cell line (3 biological replicates) was filtered to an effect size of at least 2-fold change and adjusted *p*-value of less than 0.005 with a cutoff q-value 0.05. Pathway and biological processes analysis of all DEGs was performed using Ingenuity Pathway Analysis (IPA, Qiagen, RRID:SCR_008653). DEGs were filtered to select the targets altered in both cell lines. The enrichment analysis and pathway hierarchical clustering based on overlapping DEGs were performed using REACTOME Cytoscape (RRID:SCR_003032) as described (18) (DEGs list and pathway list provided in the supplemental file). Results were visualized using GraphPad (RRID:SCR_002798).

## Statistics

Results are reported as the mean ±S.D. or mean±S.E. Two-way ANOVA approach with Tukey’s or Śidak’s post hoc analysis was used to calculate *p-*values in GraphPad Prism, version 10.0.1 (RRID:SCR_002798), *p*-values <0.05 were considered to be statistically significant.

## Data availability

The pathway enrichment analysis was performed using the data obtained from Gene Expression Omnibus (GEO) at <u>GSE67051,</u> and analyzed data are available in the supplemental files. Other data generated in this study are available upon request to the corresponding author.

## RESULTS

### Erlotinib therapy attenuates fructose-2,6-bisphosphate-dependent glucose metabolism in PC9 and HCC827 cells

Despite high sensitivity to EGFR-targeting therapies, certain mutEGFR cells manage to persist during the initial treatment, thereby contributing to the rise of resistant cell populations (19). To gain insight into the molecular perturbations driven by erlotinib therapy, we performed gene expression profiling of PC9 and HCC827 cells exposed to erlotinib (2 µM for 8 days) using publicly available data (dataset GSE67051 (17)). Using a q-value cutoff ≤ 0.05 with |log_2_FC| ≥2, we identified 2099 and 1198 differentially expressed genes (DEGs) upon exposure to erlotinib in PC9 and HCC827 cells, respectively. DEGs were filtered to select the targets altered in both cell lines and profiled using Ingenuity Pathway. We found “glucose metabolism “, “cell cycle”, and “DNA replication, recombination, and repair” among the pathways significantly changed upon erlotinib treatment (Fig. 1A, D). Notably, the “glucose metabolism” node had a negative activation z-score suggesting attenuated glucose metabolism in response to erlotinib in both cell lines. Interestingly, hierarchical clustering based on DEGs revealed significant changes in the specific branch of carbohydrate metabolism associated with the regulation of glycolysis by F2,6BP (p-value 0.032; Suppl.File1).

**Fig. 1.**
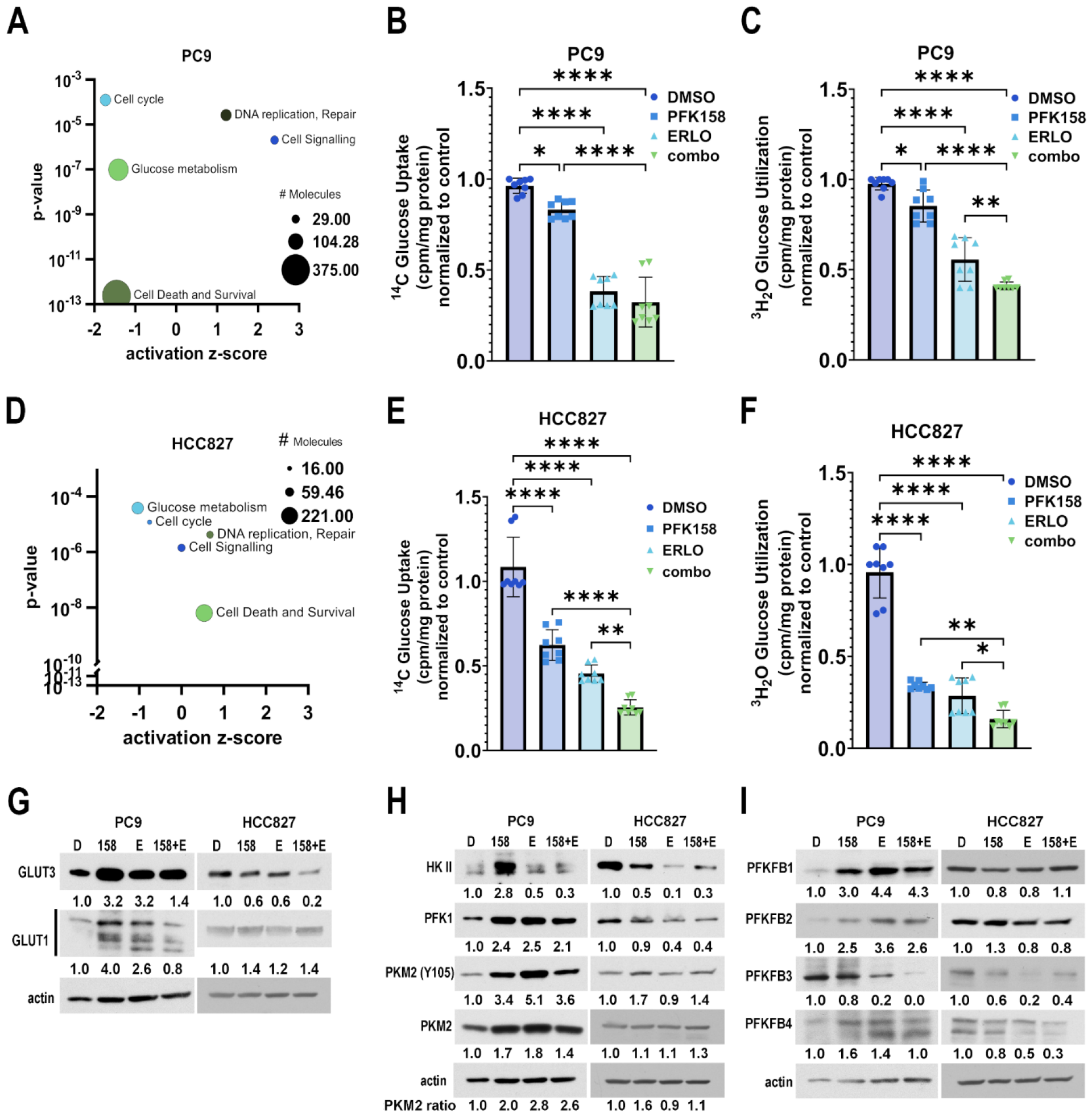
PFKFB3 maintains glycolytic flux in lung cancer cells exposed to erlotinib. Pathways enriched in response to erlotinib treatment (2 µM, 8 days) in PC9 (**A**) and HCC827 (**D**) cells. Data represent the number of differently expressed genes (DEG) for each pathway, *p*-value, and activation z-score compared to vehicle-treated cells (DMSO) (n=3). PC9 and HCC827 cells were exposed to the indicated treatments for 24h. ^14^C glucose uptake was measured in PC9 (**B**) and HCC827 (**E**) cells and normalized to the vehicle (DMSO). Individual data points represent 1 biological replicates (n=6-8). Glycolysis, measured as ^3^H_2_0 release by enolase in PC9 (**C**) and HCC827 (**F**) cells, was normalized to the vehicle (DMSO). Individual data points represent 1 biological replicate (n=8). **G-I**, metabolic enzyme protein expression in whole cell lysates assessed in immunoblotting. β-actin was used as a loading control. Target/actin ratios were quantified using densitometry and normalized to vehicle-treated samples (D). PKM2 phosphorylation ratio was calculated by dividing PKM2 (Y105) by the total PKM2 for each treatment and normalizing to vehicle-treated cells. **B,C,E,F**: Statistical analysis by ONE-WAY ANOVA with Tukey’s *post hoc* tests (*p*-values are shown as follows: *, <0.05; **, <0.01; and ****, <0.0001.) D - DMSO, 158 – PFK-158, E – erlotinib, 158+E - PFK-158 plus erlotinib.

### PFKFB3 maintains glycolytic flux in PC9 and HCC827 cells exposed to erlotinib

We previously showed that PFKFB3, a key regulator of glycolytic flux, regulates F2,6BP production and controls EGFR-mediated glycolysis in NSCLCs (4). Moreover, we demonstrated that PFKFB3 supports the survival of NSCLC cells exposed to erlotinib. Based on our previous work, we hypothesized that PFKFB3 maintains glycolytic flux in cells under erlotinib therapy. To dissect the role of PFKFB3 in NSCLC cells subjected to erlotinib, we inhibited PFKFB3 function with the small molecule inhibitor PFK-158, previously described by our group (20), and examined glucose uptake and utilization. PFKFB3 or EGFR inhibition alone significantly decreased the uptake of radiolabeled 2-[^14^C]-deoxyglucose in both cell lines (Fig. 1B, E). Dual therapy reduced glucose influx by 68% and 78% in PC9 and HCC827 cells, respectively, when compared to vehicle-treated cells. Next, we evaluated the glycolytic flux in these cells by measuring the release of ^3^H_2_O from 5-[^3^H]-glucose via enolase, which is downstream of PFKFB3-regulated PFK-1 in the glycolytic pathway. Erlotinib or PFK158 treatment significantly inhibited ^3^H_2_O release, indicating reduced glycolysis in both cell lines (Fig. 1C, F). Exposure to PFK-158 further decreased glycolytic flux in erlotinib-treated cells, resulting in a 60% (PC9) and 84% (HCC827) decrease compared to vehicle-treated cells. Given that glycolytic flux is tightly regulated at many steps, we assessed the levels of key glycolytic enzymes in response to erlotinib treatment for 48 hours alone or in combination with PFK-158. First, we evaluated the expression of glucose transporters in response to individual or combined treatments. Immunoblotting revealed an increased expression of glucose transporters GLUT1 and GLUT3 in PC9 cells exposed to erlotinib or PFK-158, which was blocked in the combined treatment (Fig. 1G). While treatment with erlotinib or PFK-158 moderately induced GLUT1 expression in HCC827 cells, decreased GLUT3 expression correlated with reduced glucose uptake in response to different therapies (Fig. 1E, G). Together, these data suggest that elevated expression of glucose transporters upon EGFR or PFKFB3 inhibition fails to sustain glucose uptake in PC9 and HCC827 cells.

Next, we examined the expression of the key enzymes controlling the glycolytic flux. PFKFB3 inhibition led to cell line-specific effects on hexokinase II (HKII) expression, with elevated HKII levels observed in PC9 cells and reduced expression in HCC827 cells. In line with previously published reports (21,22), exposure to erlotinib significantly reduced HKII expression, and this erlotinib-mediated effect was maintained upon dual treatment in PC9 and HCC827 cells (Fig. 1H). Decreased HKII levels correlated with reduced glycolytic flux in PC9 and HCC827 cells (Fig. 1C, F). Exposure to single or combined treatment in PC9 cells led to an increase in phosphofructokinase 1 (PFK1) expression. Interestingly, this compensatory increase in PFK1 failed to restore the glycolytic flux in PC9 cells treated with erlotinib, PFK-158 or the combination. In contrast, we observed an erlotinib-mediated decrease in PFK1 expression in HCC827 cells. Our data suggest that exposure to erlotinib constrained glycolytic flux at the HKII step, limiting the glucose metabolism within the glycolytic pathway in PC9 and HCC827 cells. Our data indicate that reduced glycolytic flux upon combined treatment results from a decrease in glucose uptake followed by limited glucose phosphorylation by HKII. Next, we assessed the glycolytic flux towards the TCA cycle by evaluating the expression of pyruvate kinase 2 (PKM2). We found that exposure to PFK-158 promoted PKM2 Y105 phosphorylation (inhibitory phosphorylation site) (23) in both cell lines, suggesting reduced glucose utilization in the TCA cycle. While exposure to erlotinib resulted in elevated Y105 PKM2 phosphorylation in PC9 cells, we found no difference in PKM2 levels in HCC827 cells. An erlotinib-mediated effect on PKM2 phosphorylation was maintained upon PFKFB3 inhibition in both cell lines. Our data suggest that PFKFB3 inhibition reduces glucose utilization in the TCA cycle in erlotinib-treated cells. Finally, to dissect the specific contribution of the PFKFB3 isoform to F2,6BP-dependent glycolysis under erlotinib therapy, we assessed the expression of all PFKFB isoforms in PC9 and HCC827 cells exposed to different treatment regimens. Consistent with our previous observations (4), EGFR inhibition reduced PFKFB3 expression by 80% in both PC9 and HCC827 cells (Fig. 1I). A compensatory increase in the expression of PFKFB1, PFKFB2 and PFKFB4 isoforms in response to EGFR or PFKFB3 inhibition in PC9 cells indicated a tight crosstalk between PFKFB isoforms to sustain glycolytic flux (Fig. 1I). Accordingly, the lack of compensatory PFKFBs expression in HCC827 cells correlated with a drastic decrease in glycolytic flux in response to EGFR or/and PFKFB3 inhibition compared to PC9 cells (Fig. 1C, F, I). The strong correlation between PFKFB3 function (expression and inhibition) and the treatment-mediated reduction in glycolytic flux in PC9 and HCC827 suggests that PFKFB3 supports glucose metabolism in cells exposed to erlotinib.

### PFKFB3 inhibition reduces glucose utilization in the glycolysis and TCA cycle, depleting ATP in the NSCLC cells exposed to erlotinib

Our findings indicating a decrease in HKII expression following erlotinib treatment suggest a rerouting of glucose utilization away from glycolysis. To evaluate the metabolic fate of glucose under EGFR or PFKFB3 inhibition, we conducted stable isotope tracer experiments using [U-^13^C]-glucose. Briefly, PC9 cells were treated with the inhibitor(s) for 12 hours, followed by supplementation with ubiquitously labeled glucose for an additional 24 hours. First, we analyzed glucose utilization in the initial steps of glycolysis by tracing the enrichment of M+6 hexose intermediates. PFKFB3 inhibition led to a 1.4-fold enrichment in M+6 sorbitol in erlotinib-treated cells, while individual treatments had no effect on isotopologue abundance (Fig. 2A). This elevated sorbitol labeling indicated a rerouting of glucose towards the polyol pathway (PP), typically activated under hyperglycemic conditions (such as excess glucose that cannot be utilized in glycolysis due to reduced HKII expression). The evolutionary conservative PP includes 2 steps: glucose conversion to sorbitol by aldose reductase (AKR1B1, rate-limiting enzyme) followed by conversion to fructose by sorbitol dehydrogenase (SDH). PFKFB3 inhibition significantly reduced M+6 fructose abundance, suggesting a reduction in PP flux (Fig. 2B). Conversely, erlotinib treatment resulted in elevated M+6 fructose labeling, indicating increased PP flux (Fig. 2B). Notably, PFK-158 treatment diminished the difference in fructose enrichment between the erlotinib- and combo-treated cells, indicating that PFKFB3 inhibition reduced PP flux in erlotinib-treated cells. This reduction in M+6 fructose abundance correlated with elevated M+6 sorbitol labeling, further indicating reduced PP flux towards fructose synthesis. To comprehensively assess the PP flux, we compared total sorbitol and fructose levels between treatment regimens (Fig. 2A, B, solid lines). PFK-158-induced sorbitol accumulation coincided with decreased fructose levels in erlotinib-treated cells, confirming reduced PP flux.

**Fig. 2.**
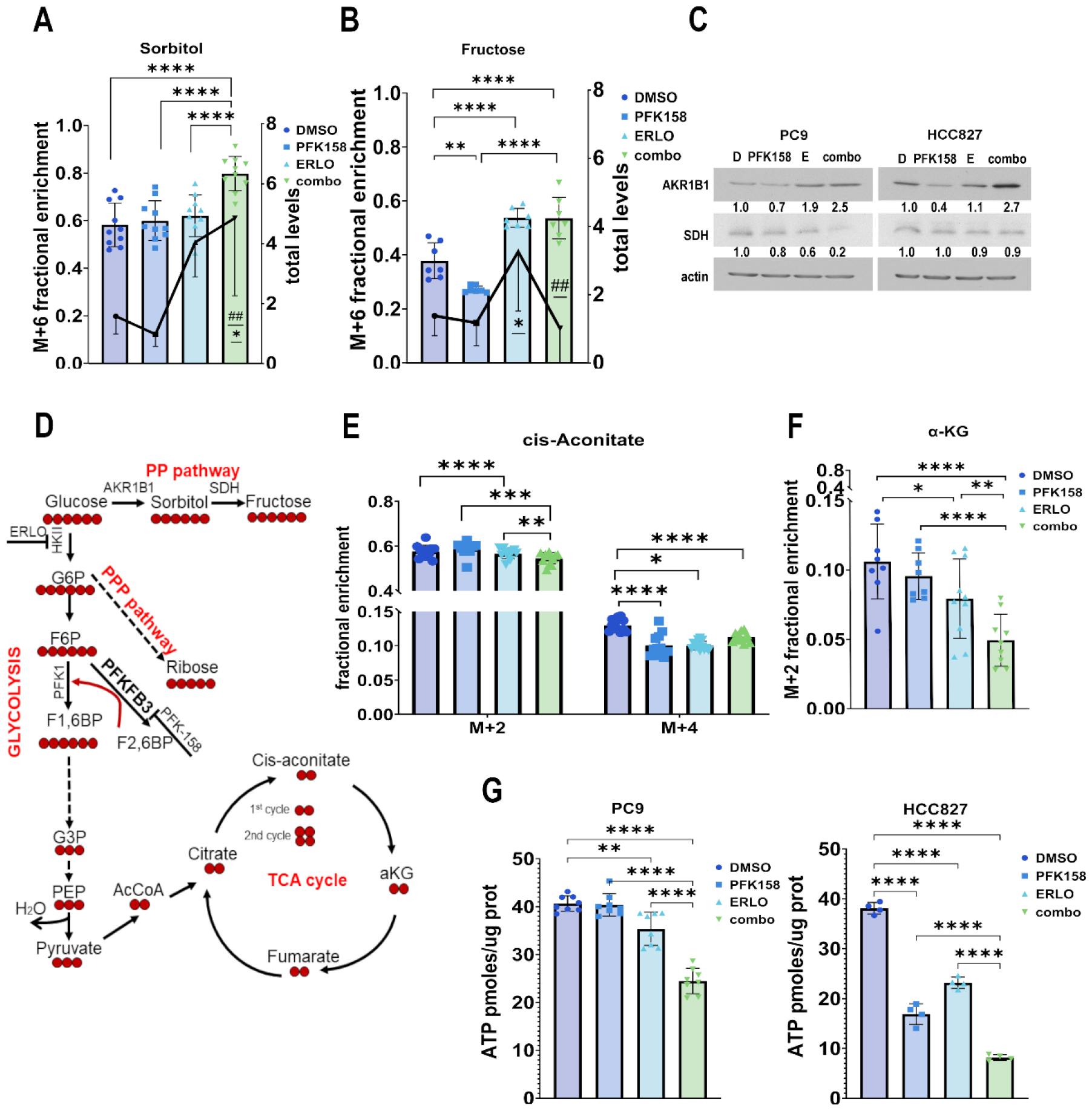
PFKFB3 inhibition reduces glucose utilization in the glycolysis and TCA cycle, depleting ATP in erlotinib-treated cells. PC9 cells were exposed to the indicated treatments for 36 hours (including 24h incubation with tracer - [U-13C]-glucose). Fractional enrichment of fully labeled sorbitol (M+6, **A, left axis**) and fructose (M+6, **B, left axis**); total levels of sorbitol (**A, right axis**) and fructose (**B, right axis**). Total levels were normalized to vehicle-treated samples. Data presented from 2 independent experiments with individual data points representing 1 biological replicate (n=7-10). Statistical analysis by TWO-WAY ANOVA with Tukey’s *post hoc* tests (*p*-values are shown as follows: *, <0.05; **, <0.01; and ****, <0.0001.) (*p*-values ##, <0.01 when compared to ERLO). **C**, AKR1B1, and SDH expression in PC9 and HCC827 whole cell lysates. β-actin was used as a loading control. Target/actin ratios were quantified using densitometry and normalized to vehicle-treated samples (D). **D**, Scheme of [U-13C]-glucose utilization in glycolysis and the TCA cycle. Fractional enrichment of cis-aconitate (**E**) and a-KG (**F**)(n=8). Statistical analysis by TWO-WAY ANOVA with Tukey’s *post hoc* tests (*p*-values: *, <0.05; **, <0.01; ***, <0.001; ****, <0.0001). **G**, total ATP levels (in pmol/µg protein) in PC9 and HCC827 cells exposed to different treatments. Data presented from 3 independent experiments (n=8) Statistical analysis by ONE-WAY ANOVA with Tukey’s *post hoc* tests (*p*-values: **, <0.01; and ****, <0.0001). Aldo-keto reductase family 1 member B (AKR1B1), sorbitol dehydrogenase (SDH), alpha-ketoglutarate (a-KG), DMSO (D), erlotinib (E, erlo), combo (PFK-158 plus erlotinib).

To elucidate the molecular mechanism underlying altered PP flux, we evaluated the expression of AKR1B1 and SDH enzymes. Importantly, PFKFB3 inhibition attenuated AKR1B1 expression in both cell lines (Fig. 2C). Consistent with previously published data (19), exposure to erlotinib upregulated AKR1B1 expression in PC9 cells but had no effect in HCC827 cells (Fig. 2C). As expected, the combination therapy led to a 2.5-fold increase in AKR1B1 expression, confirming elevated sorbitol synthesis in PC9 and HCC827 cells. While individual treatments moderately inhibited SDH expression in PC9 cells, combination therapy reduced SDH expression by 80%. Consequently, we observed an accumulation of sorbitol in erlotinib-treated cells upon PFKFB3 inhibition, further suggesting halted PP flux. Conversely, we observed insignificant changes in SDH expression in HCC827 cells. Additionally, there were no significant changes in the abundance of other M+6 hexose intermediates within the glycolytic pathway. Given that the labeling of hexoses within glycolysis was not the primary focus of this study, the labeling timing was adapted to track glucose utilization in tangential pathways (PP and PPP) and the TCA cycle. Therefore, the lack of significant changes in the abundance of M+6 glucose-6-phosphate or fructose-6-phosphate isotopologues can be attributed to our specific labeling approach. Taken together, our data suggest that PFKFB3 sustains glycolytic flux in erlotinib-treated cells by supporting glucose utilization within the polyol pathway.

To investigate the impact of dual therapy on glucose utilization within the TCA cycle, we evaluated the fractional enrichment of M+2 and M+4 isotopologues (1^st^ and 2^nd^ runs of the TCA cycle, respectively) of TCA metabolites. We observed reduced glucose carbon incorporation in M+4 cis-aconitate upon PFK-158 treatment, consistent with decreased glycolytic flux (Fig. 2D). Additionally, erlotinib treatment led to decreased M+2 and M+4 isotopologue labeling of several TCA intermediates, including cis-aconitate and α-ketoglutarate (Fig. 2D, E). Importantly, inhibition of PFKFB3 significantly reduced the abundance of these M+2 isotopologues in erlotinib-treated cells (Fig. 2D, E). The different labeling dynamics observed in cells exposed to PFK-158 (saturated M+2 labeling coinciding with reduced M+4 cis-aconitate levels) and erlotinib (reduced M+2 labeling) compared to vehicle-treated cells correlated with staggered glycolytic flux (Fig. 1C), further suggesting the contribution of glycolysis on the efficacy of TCA cycle.

Given that glucose metabolism plays a fundamental role in energy homeostasis, we next sought to assess the effect of attenuated glucose metabolism on ATP production in erlotinib-treated cells. Exposure to PFK-158 attenuated ATP production in HCC827 cells while having no significant effect in PC9 cells. In line with previous findings (22), we observed a significant decrease in ATP production in response to erlotinib in both cell lines (Fig. 2F). Importantly, PFKFB3 inhibition in erlotinib-treated cells caused a dramatic reduction in ATP production in PC9 (31%) and HCC827 (65%) cells when compared to erlotinib-treated cells. Notably, changes in ATP levels upon treatment correlated with the levels of glycolysis in both cell lines. These data confirm that cells under EGFRi therapy rely on glycolysis to maintain ATP production. Also, PFKFB3 inhibition in the cells under EGFRi therapy effectively reduced glucose utilization within glycolysis and the TCA cycle, resulting in ATP depletion.

### PFKFB3 inhibition mitigates redox capacity of cells during erlotinib therapy

Glycolysis, polyol pathway, and the TCA cycle play fundamental roles in maintaining redox homeostasis (24,25). Building on this and given the role of PFKFB3 in regulating oxidative stress homeostasis (26,27), we hypothesized that PFKFB3 inhibition would disrupt redox homeostasis in erlotinib-treated cells. First, we assessed oxidative stress levels by measuring reactive oxygen species (ROS, i.e., hydroxyl radical OH levels) in PC9 and HCC827 cells exposed to different treatment regimens. As expected, PFKFB3 inhibition triggered dramatic ROS accumulation in both cell lines (Fig. 3A). Simultaneously, exposure to erlotinib resulted in elevated ROS in HCC827 cells while having no effect on PC9 cells. Notably, PFKFB3 inhibition promoted oxidative stress in PC9 and HCC827 cells under erlotinib therapy. Recent studies have indicated that erlotinib-treated cells exhibit an increased redox capacity to sustain survival during therapy (19,28). Given that PFK-158 induces oxidative stress, we hypothesized that PFKFB3 inhibition could potentially overcome the redox capacity of NSCLC cells under EGFR therapy. Recently, it has been shown that the elevated redox capacity of the TKI-resistant cells is supported by glutathione peroxidases (GPX), including GPX4, which catalyzes the reduction of peroxides via oxidation of reduced glutathione (GSH) (19,29). We assessed GPX4 levels and found that erlotinib treatment stimulated GPX4 expression in both cell lines (Fig. 3B). Importantly, we found that PFKFB3 inhibition decreased GPX4 expression as a single therapy and attenuated the erlotinib-driven effect on GPX4 expression in both cell lines. Elevated GSH oxidation correlated with PFK158-driven ROS levels and erlotinib-driven GPX4 expression in both cell lines, confirming therapy-driven oxidative stress (Suppl. Fig1S).

**Fig. 3.**
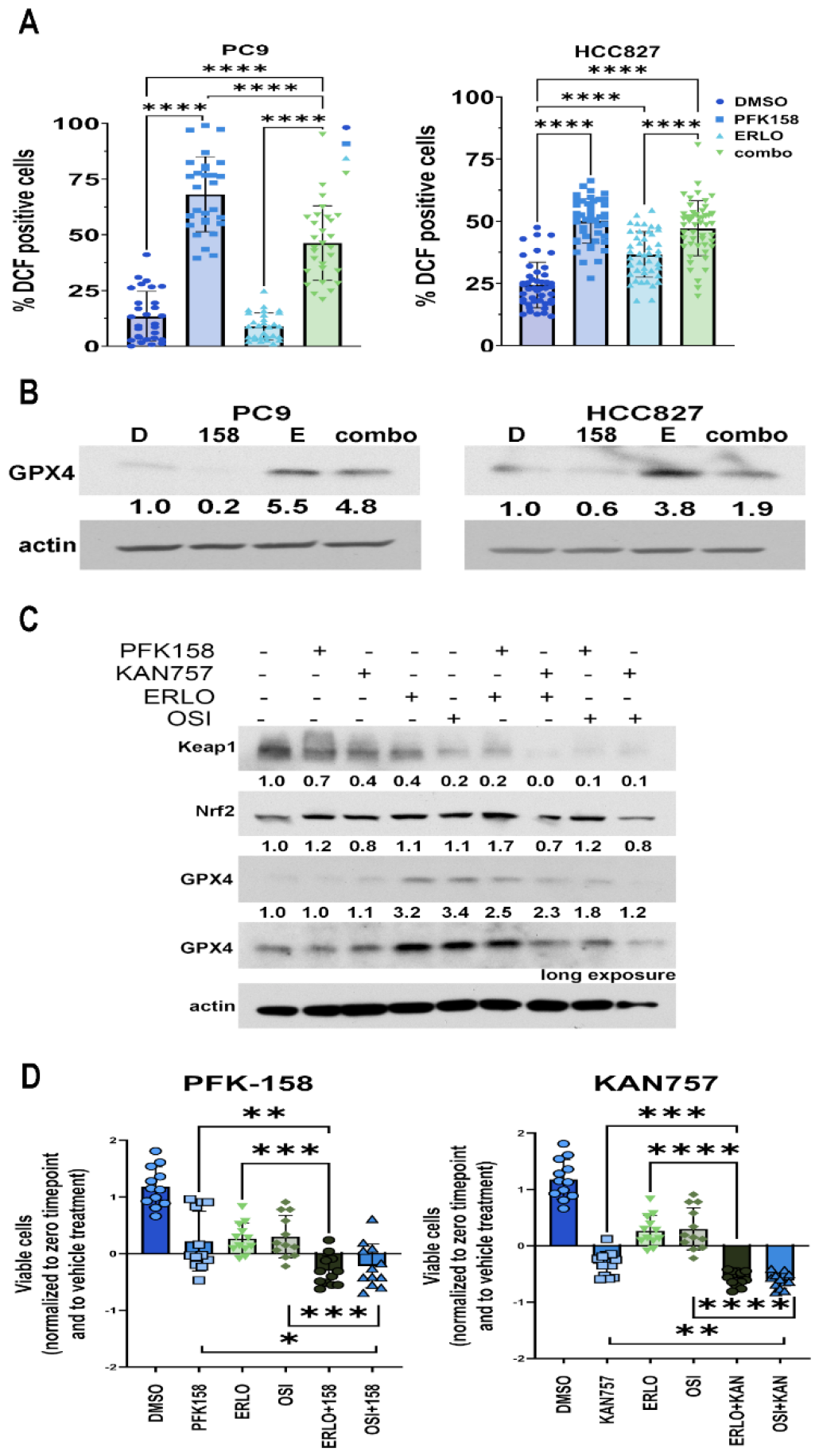
PFKFB3 inhibition mitigates the redox capacity of cells during erlotinib therapy. NSCLCs were exposed to the indicated treatments for 24h. **A**, Cellular ROS levels in individual PC9 and HCC827 cells were assessed by measuring the oxidation of 2’,7’-dichlorofluorescein diacetate (DCFDA) to 2’,7’-dichlorofluorescein (DCF). Cells were analyzed using the CellProfiler pipeline, and the number of DCF-positive cells was normalized to the total number of cells per field for each treatment condition. Data presented from 3 independent experiments. Individual data points represent 1 field (number of fields, PC9 n=29, HCC827 n=45). **B**, GPX4 expression in PC9 and HCC827 whole cell lysates. **C**, Representative immunoblot of Keap1, Nrf2, GPX4 expression in PC9 whole cell lysates. β-actin was used as a loading control. Target/actin ratios were quantified using densitometry and normalized to vehicle-treated samples. **D**, Viability of PC9 cells in response to different treatments was evaluated by trypan blue exclusion. Shown are changes in the numbers of viable cells between 0 and 24 hours post-treatment, normalized to vehicle-treated cells (DMSO). Individual data points represent 1 biological replicate from 3 independent experiments (n=12). **A, D**, Statistical analysis by ONE-WAY ANOVA with Tukey’s *post hoc* tests (*p-*values are shown as follows: *, <0.05; **, <0.01; ***, <0.001, and ****, <0.0001). Glutathione peroxidase 4 (GPX4), Kelch-like ECH-associated protein 1 (Keap1), nuclear factor erythroid 2–related factor 2 (Nrf2), DMSO (D), PFK-158 (158), erlotinib (E, erlo), combo (PFK-158 plus erlotinib), EGFR inhibitor osimertinib (OSI), PFKFB3 inhibitor KAN0438757 (KAN).

To validate that the effect of PFKFB3 inhibition on GPX4 expression is target-specific, we exposed PC9 cells to IC_50_ concentrations of two different PFKFB3 inhibitors (PFKFB3i, PFK-158 (7,5 µM) or KAN0438757 (30) (KAN757, 15 µM, as established in drug titration experiments, see Fig3 Suppl.) either alone or in combination with 1 µM EGFR inhibitors (EGFRi, erlotinib or osimertinib) for 24 hours. The ROS-mediated cellular response to oxidative stress was assessed based on the activation of the antioxidant Keap-Nrf2 pathway (31). Exposure to either PFKFB3i or EGFRi alone induced the degradation of the negative redox-sensor protein Keap1 in PC9 cells. Importantly, we observed a near complete loss of Keap1 in response to combined treatment, indicating significant activation of antioxidant signaling (Fig. 3C). While PFK-158 or EGFRi had no effect on the expression of transcription factor Nrf2, KAN757 moderately attenuated Nrf2 levels. Next, we assessed GPX4 expression and found that EGFRi-driven GPX4 expression was diminished by PFKFB3 inhibition in all combined treatments (Fig. 3C.) These data suggest that PFKFB3 inhibition disrupts the redox capacity of NSCLC cells under erlotinib therapy by downregulating GPX4 expression and limiting the antioxidant response. Given that elevated redox homeostasis supports the survival of the cells under erlotinib therapy (19), we evaluated the effect of PFKFB3 inhibition on the viability of the cells exposed to erlotinib. PFKFB3 inhibition with PFK-158 (Fig. 3D, left panel) or KAN757 (Fig. 3D, right panel) significantly attenuated the viability of the cells exposed to EGFRi (complete statistical data available in Suppl. Table 1S). Our findings indicate that PFKFB3 plays a crucial role in supporting redox homeostasis in lung cancer cells, thereby enabling their survival during EGFRi-therapy.

### PFKFB3 inhibition triggers oxidative DNA damage in TKI-treated cells

To gain insight into the molecular mechanism contributing to the diminished cell survival of erlotinib-treated cells upon PFKFB3 inhibition, we aimed to investigate the effect of PFKFB3-mediated oxidative stress on cell homeostasis. Given that elevated sorbitol and ROS trigger oxidation of cellular macromolecules, including DNA (6,32), we sought to assess the effect of combined therapy on DNA oxidation in PC9 and HCC827 cells. Given that guanine has the lowest redox potential among DNA bases (33), we initially evaluated the levels of oxidized guanine (8-oxo-G) in the cells in response to different treatment regimens. Immunocytochemistry revealed a significant accumulation of 8-oxo-G in a PFK-158-dependent manner in PC9 and HCC827 cells (Fig. 4A, B). Considering that elevated DNA oxidation can result from reduced DNA repair capacity and that PFKFB3 can directly affect DNA repair (9,30), we hypothesized that PFKFB3 is involved in the DNA damage response or repair in lung cancer cells insulted by ROS. Under standard conditions, oxidized nucleotides can be directly removed from DNA via the base-excision repair (BER) mechanism. Base excision repair requires the recruitment of the appropriate DNA glycosylases to the chromatin, where the damaged base will be removed (34). To dissect the role of PFKFB3 in ROS-driven DNA damage repair, we silenced PFKFB3 and assessed the expression of different BER targets upon erlotinib treatment. Immunoblotting revealed that PFKFB3 silencing dramatically decreased the expression of DNA-glycosylases MPG, UNG1 and 2, and NTHL1 (Fig. 4C.) The described DNA glycosylases are responsible for recognizing and removing oxidized, alkalized, or methylated nucleotides at the early steps of BER. Treatment with erlotinib significantly reduced levels of BER targets, while PFKFB3 silencing enhanced the erlotinib effect, resulting in near complete loss of UNG1/2 and NTHL1. Given that erlotinib decreases PFKFB3 expression, our data suggests that decreased expression of the listed targets upon erlotinib treatment is PFKFB3-mediated. The described DNA glycosylases are responsible for recognizing and removing oxidized, alkalized, or methylated nucleotides at the early steps of BER. BER type of DNA repair is usually activated in response to nucleotide modifications caused by alkylating agents or ROS produced by oxidative stress, further suggesting that the indicated DNA damage is ROS-driven.

**Fig. 4.**
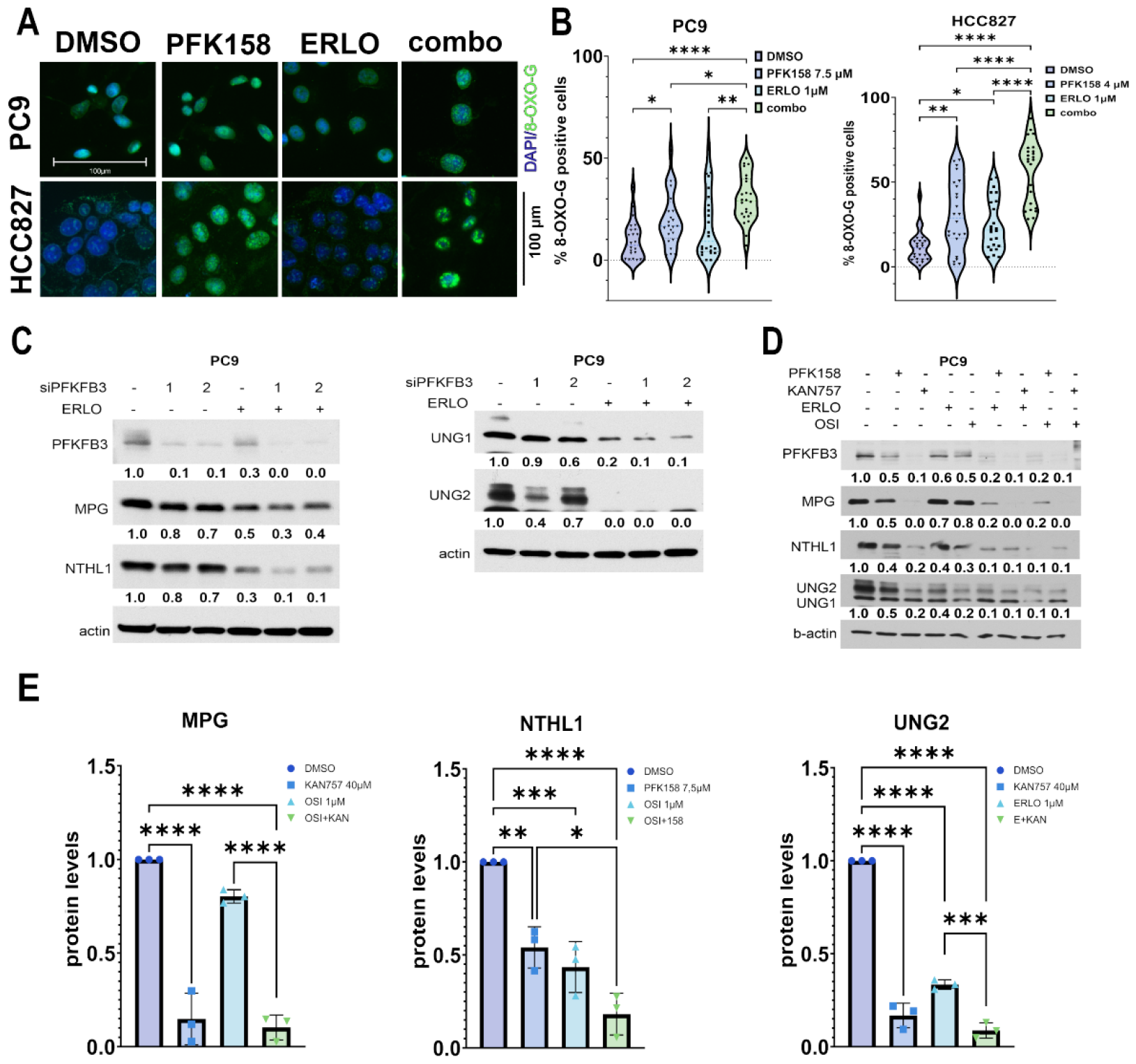
PFKFB3 inhibition triggers oxidative DNA damage in TKI-treated cells. **A**, NSCLCs were exposed to the indicated treatments for 24h. Immunocytochemical staining of oxidized guanine (8-OXO-G, green) and nuclear staining with DAPI (blue). Scale bar 100 µm. **B**, Cells were analyzed using the CellProfiler pipeline; nuclei with 8-OXO-G punctate were normalized to the total nuclei amount in the field and presented in % (n=45). **C**, PC9 cells were transfected with PFKFB3 siRNAs (si#1, si#2) followed by treatment with either vehicle or erlotinib (PC9, 0.5 μM) for 24 h. Whole cell lysates were analyzed by Western blotting with the indicated antibodies. β-actin was used as a loading control. Target ratios were quantified using densitometry and normalized to vehicle-treated samples. **D**, HCC827 cells were exposed to PFK-158 alone or in combination with antioxidant N-acetylcysteine (NAC, 5mM) for 24h. Chromatin fractions were analyzed in immunoblotting. Histone H3 was used as a loading control. **E**, PC9 cells were exposed to PFKFB3 inhibitors (PFK158, KAN757) or/and EGFR inhibitors (erlotinib, osimertinib) for 24 h. Whole cell lysates were analyzed by Western blotting with the indicated antibodies. ^β^-actin was used as a loading control. Target/loading control ratios were quantified using densitometry and normalized to vehicle-treated samples. **F**, MPG, NTHL1, and UNG2 protein levels in PC9 cells exposed to appropriate treatments were analyzed in immunoblotting in 3 independent experiments (n=3). **B, F**, Statistical analysis by ONE-WAY ANOVA with Tukey’s *post hoc* tests (*p*-values are shown as follows: *, <0.05; **, <0.01; ***, <0.001; ****, <0.0001). N-methylpurine DNA Glycosylase (MPG), Nth Like DNA Glycosylase 1 (NTHL1), Uracil-DNA glycosylase (UNG).

To further confirm that PFK-158-mediated DNA oxidation is a result of reduced BER in erlotinib-treated cells, we evaluated levels of BER targets in PC9 cells exposed to PFKFB3i alone or in combination with EGFRi for 24 hours. As expected, PFKFB3 inhibition dramatically decreased the expression of all the targets, with the KAN757 showing a more robust effect due to a much higher IC_50_ concentration when compared to PFK-158 (15 µM versus 7.5 µM, Fig. 4D.) Reduced expression of MPG, NTHL1, and UNGs in the cells exposed to erlotinib or osimertinib correlated with attenuated expression of PFKFB3. Combined therapy resulted in nearly complete loss of MPG, NTHL1, and UNG2 in PC9 cells (Fig. 4E). Our results suggest that the PFKFB3 asserts control on DNA oxidation by supporting the expression of DNA-glycosylases involved in BER.

### PFK-158 triggers DNA damage due to limited ATM-driven DDR in erlotinib-treated cells

Our data showed that inhibiting PFKFB3 triggers DNA oxidation and limits its repair through BER, suggesting elevated levels of DNA damage. Given that oxidative stress and unresolved BER can generate complex DNA lesions with double-strand breaks (DSB) (35,36), we sought to assess the levels of DNA DSB in response to PFK-158 treatment. Neutral COMET assay revealed a significant accumulation of DNA DSBs upon PFKFB3 inhibition in PC9 and HCC827 cells (Fig. 5A, B). PFK-158-mediated DNA damage was maintained in combined treatments in both cell lines. DSBs, when occur, are recognized by DNA damage response (DDR) cascade, orchestrated by two important enzymes: ATR and ATM kinases. Given that oxidative stress directly regulates the ATM function (37), we analyzed ATM activation in response to individual or combined treatment. Elevated ATM S1981 phosphorylation confirmed ATM activation upon PFKFB3 or EGFR inhibition in PC9 and HCC827 cells (Fig. 5C). Unexpectedly, we found that PFKFB3 inhibition dramatically reduced total ATM expression in both cell lines. Complete loss of ATM in erlotinib-treated cells with silenced PFKFB3 further confirmed PFKFB3-dependent ATM expression in PC9 cells (Suppl. Fig. 2S). To assess the ability of reduced ATM expression to properly activate DDR, we evaluated the phosphorylation of a direct downstream target of ATM – histone γH_2_AX (38,39) in immunocytochemistry. Elevated γH_2_AX phosphorylation at serine 139 coincided with γH_2_AX-positive foci accumulation in response to PFK-158 in PC9 and HCC827 nuclei confirmed activated DDR (Fig. 5D, E). In line with previously published data (40), exposure to erlotinib reduced S139 γH_2_AX focal accumulation by 50% compared to control treatment in both cell lines. Dual therapy marginally induced S139 γH_2_AX-foci levels in HCC827 cells while having no effect on γH_2_AX in PC9 cells compared to erlotinib treatment.

**Fig. 5.**
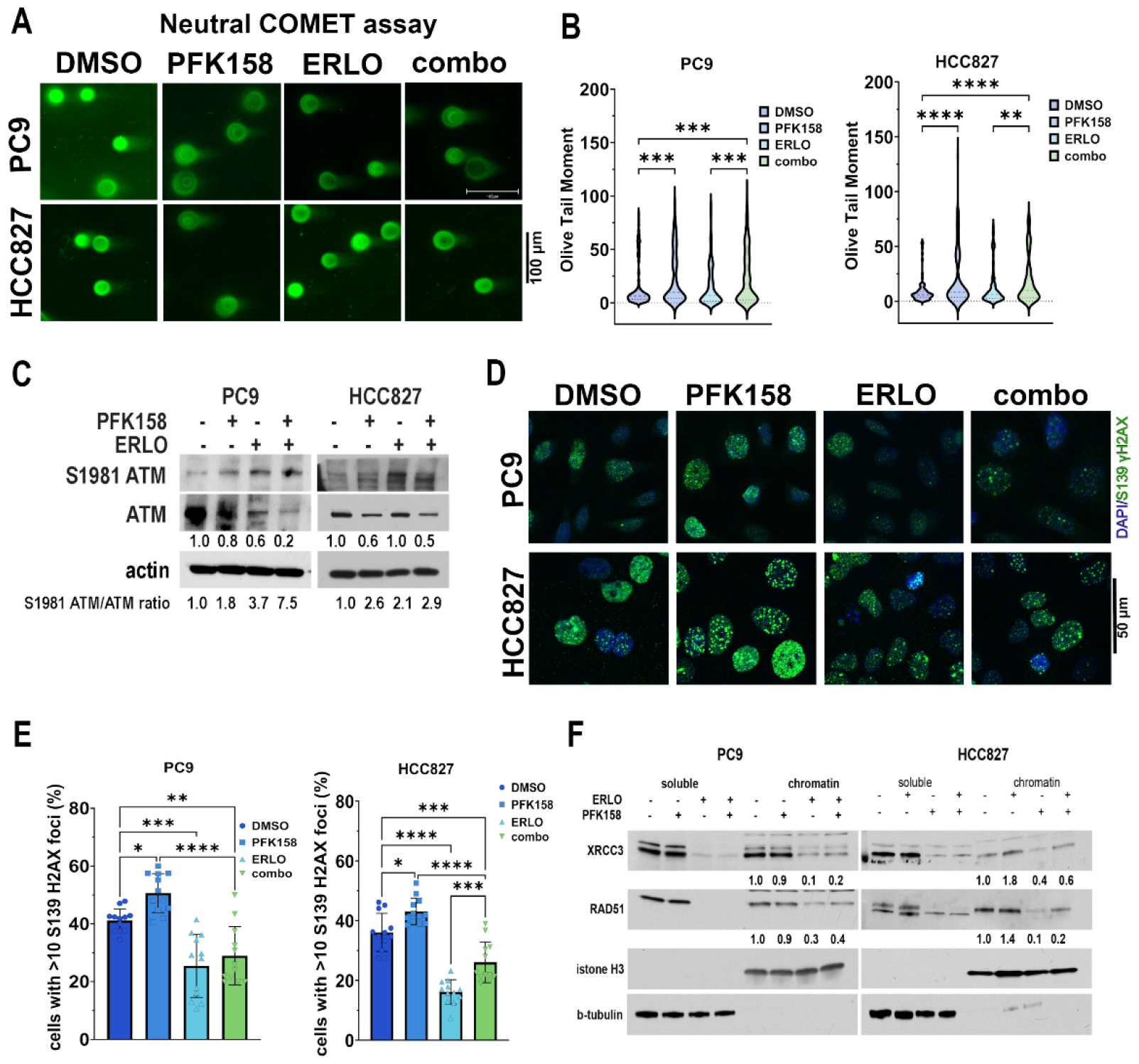
PFK-158 triggers DNA damage due to limited ATM-driven DDR in erlotinib-treated cells. **A**, NSCLCs were exposed to appropriate treatments for 24h. Visualization of a neutral comet assay showing therapy-induced DNA double-strand breaks compared to DMSO vehicle control. Scale bar 100 µm. **B**, Quantification of the neutral comet assay was performed using the OpenComet plugin for ImageJ software, and DNA breaks presented as DNA olive tail moment. The violin plot represents the data for individual nuclei from 2 independent experiments with technical replicates (n=250). **C**, ATM phosphorylation (S1981) and total protein levels in PC9 and HCC827 cells in response to treatments assessed in immunoblotting. β-actin was used as a loading control. Target/loading control ratios were quantified using densitometry and normalized to vehicle-treated samples. ATM phosphorylation ratio was calculated by dividing ATM (S1981) by the total ATM densitometry signals for each treatment and normalizing to vehicle-treated cells. **D**, Representative images of nuclei showing phosphorylated γ-H2AX foci (S139, green) and nuclear DNA stained with DAPI (blue). Scale bar 50 µm. **E**, Cells were analyzed using CellProfiler pipeline, and relative amounts of the cells with >10 γ-H2AX foci normalized to the total amount of cells per field and presented in %. Each data point represents one field analyzed (3 independent experiments, n=12). **F**, PC9 and HCC827 cells were exposed to appropriate treatment for 24h. Cytosolic (soluble) and chromatin fractions were separated and analyzed in immunoblotting. Histone H3 was used as a loading control for chromatin fraction, and β-tubulin – for soluble fraction. Target ratios were quantified using densitometry and normalized to vehicle-treated samples. **B, E**, Statistical analysis by ONE-WAY ANOVA with Tukey’s *post hoc* tests (*p-*values are shown as follows:*, <0.05; **, <0.01; ***, <0.001; ****, <0.0001). Ataxia-telangiectasia mutated (ATM), X-Ray Repair Cross Complementing 3 (XRCC3), RAD51 Recombinase (RAD51).

Importantly, in both cell lines, the levels of γH_2_AX-positive foci in response to dual therapy were significantly lower compared to the vehicle treatment, confirming that DDR is strictly limited under erlotinib therapy (Fig. 5D, E). Given that active DDR requires proper assembling of DNA repair complexes (41,42), we assessed the recruitment of DDR scaffold proteins to chromatin in response to different treatment regimens. We found that PFKFB3 inhibition promoted the recruitment of RAD51 and XRCC3 to the chromatin in HCC827 cells while having no effect in PC9 cells (Fig. 5F). In line with previous findings (43,44), administration of erlotinib resulted in reduced RAD51 expression, as evidenced by decreased RAD51 levels in both soluble and chromatin-bound fractions. As a result, we observed 80% and 90% reduction in RAD51 presence in the chromatin fraction of PC9 and HCC827 cells, respectively. Similarly, exposure to erlotinib resulted in reduced XRCC3 expression, decreasing XRCC3 recruitment to chromatin by 60% and 70% in PC9 and HCC827 cells, respectively (Fig. 5F.) Erlotinib-mediated effect on scaffold proteins expression and chromatin recruitment was maintained upon combined treatment in both cell lines. Our results indicate that erlotinib-treated cells have limited DDR as established by reduced S139 γH_2_AX focal accumulation coincided and attenuated expression of DDR scaffold proteins. These observations align with previous findings (45,46), showing that cells with ATP depletion and limited metabolic flexibility (i.e., inability to stimulate glycolysis, see Fig. 2) cannot sustain DDR. Altogether, these results suggest that PFK-158-driven DNA oxidation is translated into unresolved double-strand DNA breaks due to suspended DDR in erlotinib-stressed lung cancer cells.

### Reduced *de novo* nucleotide synthesis and oxidative stress contribute to cell death in NSCLCs under EGFRi therapy

Elevated ROS can cause the oxidation of the intracellular nucleotides, depleting the cellular nucleotide pool (47). Impaired DDR favors glucose rerouting to PPP for *de novo* nucleotide synthesis and NADPH production to counteract oxidant stress (48). The DDR sensor ATM controls PPP by directly regulating the activity of glucose-6-phosphate dehydrogenase (G6PD), promoting the *de novo* synthesis of nucleotides required for DNA repair (48,49). G6PD, a rate-limiting enzyme in PPP, is upregulated in NSCLC cells with disturbed redox states, contributing to TKI-resistance (50,51). Finally, reduced *de novo* nucleotide synthesis hampers DNA repair (52). Given that PPP supports redox homeostasis and DNA repair, we sought to assess the levels of *de novo* synthesized nucleotides available in the cells under different treatment regimens. Building on our previous findings that erlotinib rerouted glucose utilization from glycolysis towards the polyol pathway (Fig. 2A), we hypothesized that EGFR inhibition reduced glucose oxidation in the oxidative branch of PPP. First, we assessed *de novo* nucleotide synthesis levels by evaluating the enrichment in 5-ribose moiety from ribose-5-phosphate (M+5) in nucleotides in tracing studies using PC9 cells. As expected, erlotinib treatment inhibited glucose carbon incorporation in M+5 isotopologues, while PFKFB3 inhibition had no effect on glucose carbon incorporation in 5-CMP, guanosine, UTP, ADP nucleotide precursors (Fig. 6A, suppl. Fig. 3S). The erlotinib-mediated effect was maintained in 5-CMP labeling, while PFKFB3 inhibition moderately decreased M+5 guanosine enrichment upon combined treatment (Fig. 6A.) Immunoblotting revealed non-significant changes in G6PD expression in PC9 cells in response to individual or combined therapies (Fig. 6B.) Our data suggest that the limited metabolic flexibility caused by erlotinib therapy impairs the proper activation of the PPP oxidative branch in response to PFK-158-driven oxidative stress.

**Fig. 6.**
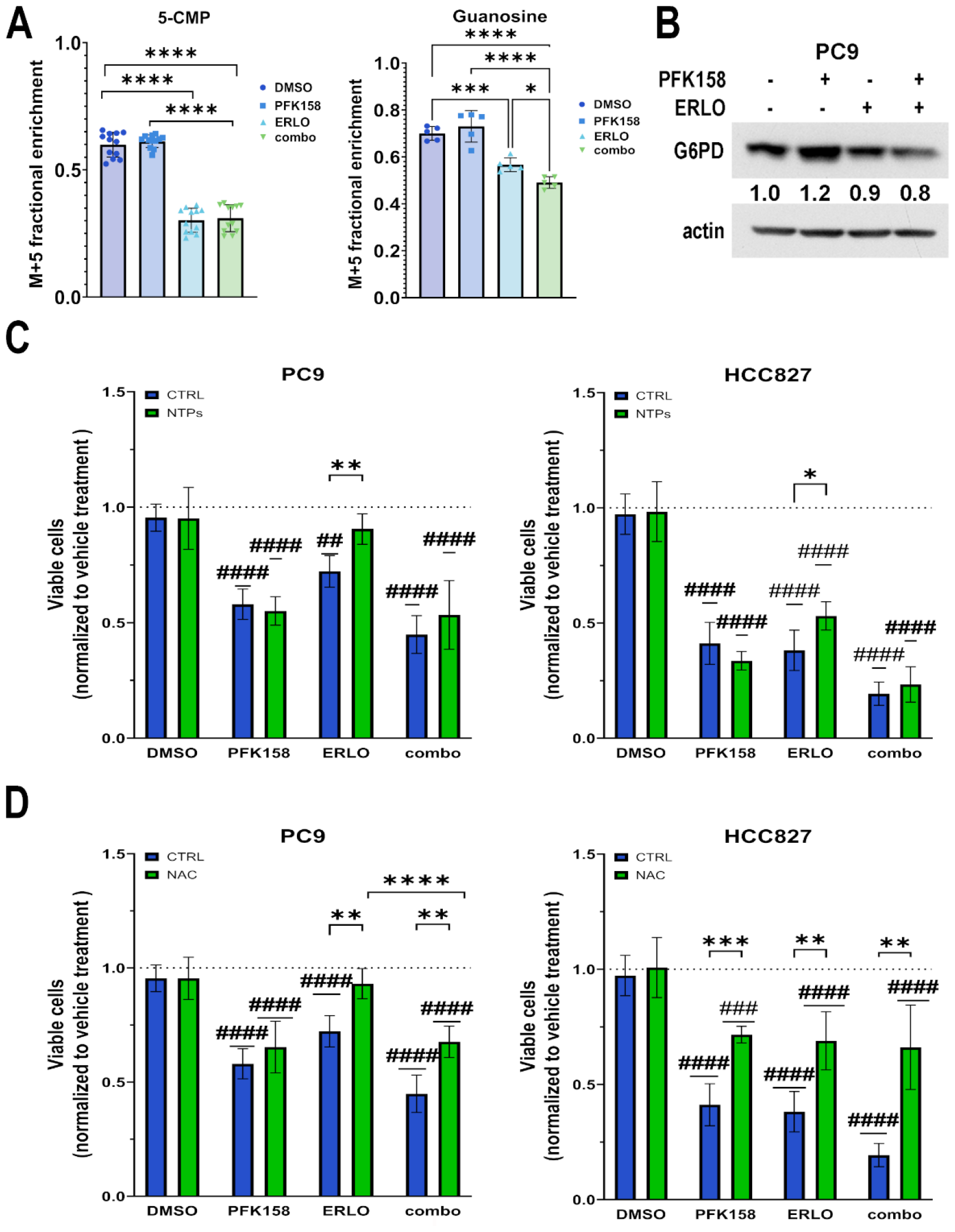
Reduced *de novo* nucleotide synthesis and oxidative stress contribute to cell death in NSCLCs under EGFRi therapy. **A**, PC9 cells were exposed to appropriate treatments for 36 hours (including 24h exposure to tracer - [U-13C]-glucose.) Fractional enrichment of M + 5 labeled nucleotides and nucleosides presented. Values represent biological replicates from 2 independent experiments (n=5-12). Data analyzed by TWO-WAY ANOVA with Tukey’s *post hoc* tests (*p*-values are shown as follows: *, <0.05; ***, <0.001; and ****, <0.0001.) **B**, PC9 cells were exposed to appropriate treatments for 24h, and whole cell lysates were analyzed in immunoblotting. Glucose-6-phosphate dehydrogenase (G6PD) expression was normalized to β-actin (loading control). Target/loading control ratios were quantified using densitometry and normalized to vehicle-treated samples. **C**, PC9 and HCC827 cells were exposed to assigned treatments for 24h with nucleotide pool replenishment in the last 6 hours. Cell viability was evaluated by trypan blue exclusion and normalized to vehicle-treated cells (DMSO). **D**, PC9 and HCC827 cells were exposed to assigned treatments for 24h in the presence of antioxidant N-acetylcysteine (NAC, 5mM, 24h). Mean ± S.E. of 3 independent experiments presented (n=12). Statistical analysis by TWO-WAY ANOVA with Tukey’s and Śidak’s *post hoc* tests (*p-*values presented as follows: ##, <0.01; ###, <0.001, and ####, <0.0001, when compared to appropriate DMSO control. When comparing NTPs or NAC effect within the same treatment regimen, *p*-values are shown as follows: *, <0.05; **, <0.01; and ****, <0.0001).

Our mass spectrometry data showed limited *de novo* nucleotide synthesis in response to erlotinib, suggesting an imbalanced nucleotide pool. We hypothesized that erlotinib-attenuated PPP, combined with elevated demand for *de novo*-synthesized nucleotides to repair PFK158-driven DNA oxidation, attenuates the viability of the cells exposed to dual therapies. Given that PFKFB3 inhibition significantly impaired BER in PC9 and HCC827 cells, we hypothesized that NTPs’ replenishment fails to improve cell viability if PFKFB3 function is inhibited. To dissect the role of NTP pool in the viability of erlotinib-treated cells, we exposed PC9 and HCC827 cells to appropriate treatments for 24 hours while replenishing the nucleotides (30 µM pool) for the last 6 hours (18-24h). In line with previous observations (53), we found that NTP supplementation significantly improved the viability of PC9 and HCC827 cells treated with erlotinib (Fig. 6C, complete statistical data available in suppl. Table 2S). At the same time, NTPs restoration failed to reverse PFK-158 effect and reinstate cell viability. Consequently, we observed no improvement in the viability of PC9 and HCC827 cells exposed to the dual therapies. These data suggest that impaired DNA repair significantly contributes to the PFK-158-mediated cytotoxic effect in erlotinib-treated cells (Fig. 3D, 6C).

Next, to confirm that PFK-158-driven cytotoxicity is ROS-mediated, we exposed PC9 and HCC827 cells to the indicated treatments in the presence of antioxidant NAC. In line with previous observations (54), NAC supplementation everted erlotinib’s effect, completely restoring cell viability in PC9 cells. However, co-treatment with NAC failed to override the PFK-158 impact, resulting in limited efficacy of ROS-scavenger in PC9 cells exposed to combined therapies (Fig. 6D). On the other hand, NAC supplementation in HCC827 cells alleviated the effect of PFK-158 and erlotinib, completely reversing the impact of individual or combined therapy (Fig. 6D). In line with previously published data (19,54-56), our results confirm that NSCLCs under erlotinib therapy rely on redox homeostasis for survival. Our findings indicate that PFKFB3 plays a multifaceted role in the survival of cells. Specifically, it aids glycolytic flux, regulates oxidative stress, and BER DNA damage response, thereby bolstering the drug tolerance of mutEGFR NSCLCs under EGFRi therapy.

## DISCUSSION

Lung cancer cells that develop tolerance to TKI therapy undergo a wide range of metabolic adaptations, including transitions to slow or non-proliferating phenotypes, switches in cell identity, and the development of immune-evasion mechanisms (57). Understanding these metabolic adaptations in cells initially tolerant to TKIs is crucial, as they can give rise to drug-resistant cell populations responsible for tumor reoccurrence (2,19,29,44). While metabolic rewiring of TKI-resistant cells has been extensively studied (58-60), the role of glycolysis in the initial response to targeted therapies remains unclear. In the current study, we demonstrate that the glycolytic regulator PFKFB3 plays a multifaceted role in the metabolic response of cells under TKI therapy. We found that EGFR inhibition significantly reduces glucose metabolism, leading to increased cell dependency on PFKFB3 to sustain glycolysis and the TCA cycle for ATP generation (Fig. 1-2). Recently, it has been demonstrated that cells surviving TKI therapy shift towards utilizing the polyol pathway for glucose utilization (19,56). This rerouting of glucose towards the PP pathway enables TKI-treated cancer cells to mitigate the negative effects of reduced HKII expression (22), which regulates glucose entry into glycolysis. Our glucose tracing studies in PC9 cells confirmed increased glucose utilization within the PP pathway in response to EGFR inhibition (Fig. 2A-B). Moreover, we observed that inhibiting PFKFB3 alone reduces the expression of AKR1B, an enzyme responsible for converting glucose to sorbitol, in PC9 and HCC827 cells. However, when combined with an EGFR inhibitor, PFKFB3 inhibition paradoxically promotes AKR1B expression. Further research is required to fully elucidate the role of PFKFB3 in the polyol pathway and its crosstalk with AKR1B1. Our data also showed that exposure to PFK-158 reduces fructose production in erlotinib-treated cells, indicating that PFKFB3 inhibition halts the PP flux. Our glucose tracing studies have also confirmed that PFKFB3 inhibition in TKI-treated cells reduces glucose utilization in the TCA cycle (see Fig. 2E-F). Considering the observed inhibitory effect of PFK-158 on the glycolytic flux, we speculate that the metabolic function of PFKFB3 becomes critical for sustaining ATP production in TKI-stressed cells (Fig. 2G). These results, along with a previously published study by our group showing that PFKFB3 inhibition limits autophagy flux and results in AMPK inactivation in erlotinib-treated cells (61) provide strong evidence that inhibiting PFKFB3 can effectively limit the metabolic adaptivity of lung cancer cells when treated with targeted therapies.

An increasing body of evidence highlights the critical role of redox capacity in the survival of stressed cells during therapy, leading to the emergence of drug-tolerant cells that persist under treatment (19). The balance between ROS production and elimination involves a complex interplay of antioxidants and redox proteins. Recently, it has been reported that elevated GPX expression supports redox homeostasis in erlotinib-treated cells (19,28,55) (Fig. 3). In our studies, we observed that PFK-158-induced oxidative stress coincided with reduced GPX4 expression. Interestingly, PFKFB3 inhibition did not lead to a proportional increase in GPX4 expression in erlotinib-treated cells, resulting in significant ROS accumulation. While the mechanism by which PFKFB3 controls GPX4 expression remains unknown, our data suggest that PFKFB3 contributes to overcoming the redox capacity of lung cancer cells by mediating GPX4 expression in TKI-treated cells. Additionally, our experiments showed that the PFKFB3 inhibitor KAN0438757 moderately decreased Nrf2 levels (Fig. 3C). While this effect could be attributed to off-target drug effects, given the high concentration applied (15 µM), we speculate that PFKFB3 might play an indirect role in the transcriptional regulation of Nrf2-dependent antioxidant response. However, this further investigation is required to confirm this speculation.

Recent studies have demonstrated the impact of cellular metabolism on DNA integrity, leading to *de novo* mutagenesis (44) and therapy resistance (62). Specifically, it has been shown that EGFR inhibition induces low-fidelity DNA polymerases and imbalanced nucleotide metabolism (44). Our findings indicate that oxidative stress driven by PFKFB3 inhibition in TKI-treated cells promotes DNA oxidation, resulting in double-strand DNA damage. While the role of PFKFB3 in regulating DNA integrity through homologous recombination (30)and miss-match repair pathways is known (30) (9), we discovered that PFKFB3 also regulates base excision repair in TKI-treated cells. Specifically, we found that PFFKB3 sustains the expression of DNA-glycosylases such as MPG, NTHL1, and UNG1/2, which recognize and eliminate modified nucleotides. The PFKFB3-dependent expression of the mitochondrial DNA-glycosylase UNG1 suggests that PFK-158-driven oxidative stress might affect mitochondrial DNA oxidation. Further investigation is required to explore the potential role of PFKFB3 in supporting mitochondrial DNA integrity, as mitochondrial DNA encodes the subunits of the oxidative phosphorylation enzyme complexes. We found that both genetic and pharmacological inhibition of PFKFB3 led to a reduction in the expression of the DNA damage sensor ATM, corroborating recent findings (6) that highlight the role of PFKFB3 in ATM-dependent DNA repair. In our studies, the DNA damage resulting from PFK-158-induced DNA oxidation was significantly exacerbated due to the overlap with limited ATM-dependent DDR from erlotinib (Fig. 5E) and ATP depletion (Fig. 2F). These align with recent research suggesting that ATP depletion activates ATM to modulate mitochondrial metabolism to sustain ATP production and Nrf1-dependent redox signaling, rather than primarily promoting DNA repair (46,63). We speculate that blocking ATM expression and ATM-driven glucose metabolism with PFK-158 results in deficient metabolism rerouting and impaired DNA repair in TKI-stressed cells.

While the precise molecular mechanism of PFKFB3-mediated ATM expression remains elusive, we speculate that PFKFB3 may orchestrate redox homeostasis and DDR by exerting control over ATM. Nevertheless, further investigation is crucial to establish the exact role of PFKFB3 in regulating ATM function, particularly in terms of its cellular localization and interaction with Nrf. The noted limitation of this study is that ATM recruitment to DNA damage sites varies between S and G1 phases, influencing the assembly of different DNA repair factors involved in repair and checkpoint activation (64). Therefore, additional investigation is needed to determine whether chromatin condensation limits the recruitment of BER targets and scaffold proteins in response to PFK-158, especially considering the durable G1 arrest promoted by EGFR inhibition in PC9 and HCC827 cells (65).

The pentose phosphate pathway plays a critical role in alleviating oxidative stress by synthesizing NADPH and ribose, essential precursors for nucleotide synthesis. The results of our study reveal that the administration of erlotinib significantly decreases the *de novo* synthesis of nucleotide precursors, with PFK-158 having a modest effect on ribose synthesis. While other studies have reported compensatory PPP activation in response to erlotinib (21) or PFKFB3 inhibition (8), we did not observe activation of the pentose-phosphate shunt with individual or combined treatment. This lack of activation could be attributed to therapy-induced glucose rerouting towards the polyol pathway (Fig. 2, 6A-C). Nucleotide supplementation reduced erlotinib’s effect on cell viability, suggesting that the NTP pool supports the viability of therapy-stressed cells. Based on our finding that PFKFB3 inhibition negates NTP supplementation’s effect on TKI-treated cells’ viability, we speculate that NTP-dependent metabolic rewiring ultimately converges to support glycolysis in therapy-tolerant cancer cells. Based on the complex role of PFKFB3 in redox homeostasis, targeting PFKFB3 emerges as a promising strategy to challenge cellular redox capacity and improve the cytotoxicity of chemotherapies in lung cancer.

Our findings underscore the importance of glycolysis in supporting redox homeostasis and DNA integrity in cells under TKI-driven stress. Based on our observations that PFKFB3 controls redox homeostasis and DDR capacities, we propose that PFK-158-driven oxidative stress and DDR-related aberrations contribute to attenuated cell viability in TKI-treated cells. Our data suggest that targeting PFKFB3, a key metabolic regulator controlling glucose utilization pathways, can limit the metabolic rewiring strategies that TKI-treated cells employ to evade cell death. We posit that aligning anti-PFKFB3 treatment with the timing of tolerance-to-resistance transition may be crucial for eradicating surviving cell populations. Administering short and precisely timed treatments could significantly improve the long-term cytotoxicity of the first-line targeted therapies.

## Supporting information

List of Antibodies

