## Supplementary material for "Role of 6-Phosphofructo-2-Kinase/Fructose-2,6-Bisphosphatase-3 in Maintaining Redox Homeostasis and DNA Repair in Non-Small Cell Lung Cancers Under EGFR-Targeting Therapy": List of Antibodies

| AB | Manufacturer | Cat No | RRID | 1' dilution |
|---|---|---|---|---|
| AKR1B1 | scbt | sc-166918 | RRID:AB_10609906 | 1/500 |
| Goat anti-mouse IgG | Thermo Fisher Scientific | 31431 | RRID:AB_10960845 | 1/10000-1/20000 |
| APNG (MPG) | scbt | sc-101237 | RRID:AB_1119016 | 1/500 |
| S1981 ATM | CST | 4526 | RRID:AB_2062663 | 1/1000 |
| ATM | CST | 2873 | RRID:AB_2062659 | 1/1000 |
| b-Tubulin | CST | 2146 | RRID:AB_2210545 | 1/1000 |
| G6PD | CST | 8866 | RRID:AB_10827744 | 1/1000 |
| GLUT1 | scbt | sc-377228 | RRID:AB_2891096 | 1/500 |
| GLUT3 | scbt | sc-74497 | RRID:AB_1124974 | 1/500 |
| Goat anti-rabbit IgG | Invitrogen | 32260 | RRID:AB_1965959 | 1/20000 |
| GPX4 | scbt | sc-166570 | RRID:AB_2112427 | 1/500 |
| HCNP (XAB2) | scbt | sc-271037 | RRID:AB_10610225 | 1/1000 |
| Histone H3 | CST | 9715 | RRID:AB_331563 | 1/1000 |
| HK I | CST | 2024 | RRID:AB_2116996 | 1/1000 |
| HK II | CST | 2867 | RRID:AB_2232946 | 1/1000 |
| Keap1 | scbt | sc-365626 | RRID:AB_10844829 | 1/500 |
| Nrf2 | scbt | sc-365949 | RRID:AB_10917561 | 1/500 |
| NTH1 (NTHL1) | scbt | sc-130644 | RRID:AB_2154558 | 1/500 |
| PFK1 | NOVUS | NBP2-75578 | RRID: AB_3095046 | 1/1000 |
| PFKFB1 | abcam | ab154573 | RRID:AB_3095047 | 1/1000 |
| PFKFB2 | CST | 13029 | RRID:AB_2232946 | 1/1000 |
| PFKFB3 | Proteintech | 13763-1-AP | RRID:AB_2162854 | 1/3500 |
| PFKFB4 | abcam | ab137785 | RRID:AB_2722775 | 1/500 |
| PKM2 | CST | 4053 | RRID:AB_1904096 | 1/1000 |
| RAD51 | CST | 8875 | RRID:AB_2721109 | 1/1000 |
| SORD (SDH) | scbt | sc-377200 | RRIB:AB_3095045 | 1/500 |
| Tyr105 PKM2 | CST | 3827 | RRID:AB_1950369 | 1/1000 |
| UDG (UNG) | scbt | sc-73639 | RRID:AB_1131097 | 1/500 |
| UNG | NOVUS | NBP1-49985 | RRID:AB_10012175 | 1/5000 |
| XRCC3 | NOVUS | NB100-165SS | RRID:AB_922791 | 1/5000 |
