## Supplemental Materials for "Role of 6-Phosphofructo-2-Kinase/Fructose-2,6-Bisphosphatase-3 in Maintaining Redox Homeostasis and DNA Repair in Non-Small Cell Lung Cancers Under EGFR-Targeting Therapy"

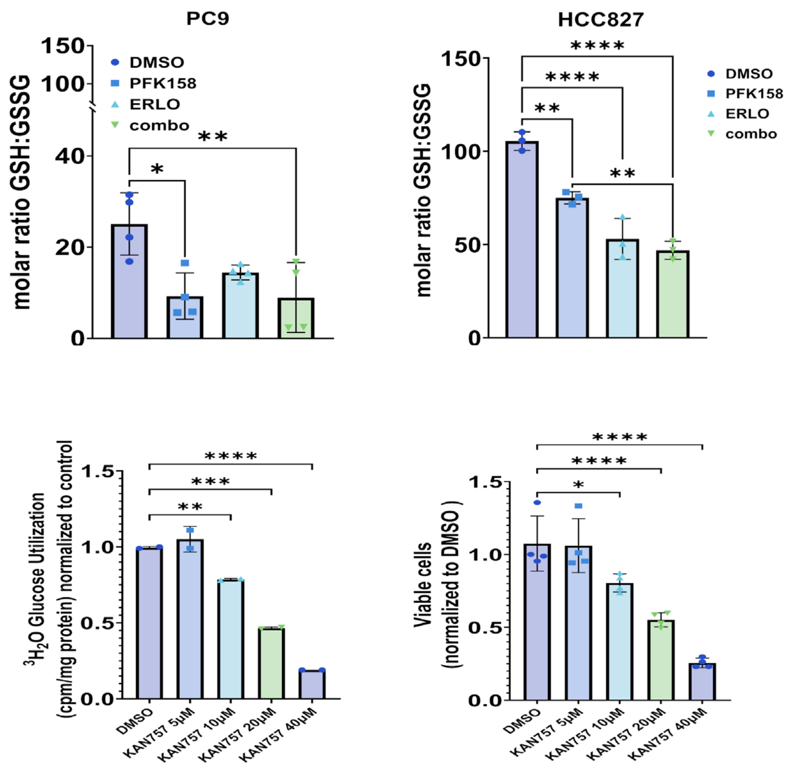

**Fig. 1S**. **TOP panel:** PC9 (left) and HCC827 (right) cells were exposed to assigned treatments for 24 h. Total, reduced (GSH), and oxidized (GSSG) glutathione levels were measured in glutathione reductase reaction. The molar ratio of GSH to GSSG is presented. Data presented from 2 independent experiments (n=4) Statistical analysis by ONE-WAY ANOVA with Tukey's *post hoc* tests (p values: *, <0.05; **, <0.01; and ****, <0.0001.)

**BOTTOM panel**: PC9 cells were exposed to different concentrations of PFKFB3 inhibitor KAN0438757 for 24h. **Left panel:** Glycolysis, measured as 3H20 release by enolase, was normalized to the vehicle (DMSO) Individual data points represent 1 biological replicate (n=4). **Right panel:** Cell survival was evaluated by trypan blue exclusion. Shown are changes in viable cell amounts when normalized to DMSO. Error bars, mean ± S.D. of two independent experiments (n = 4). ) Statistical analysis by t-test (p values: *, <0.05; **, <0.01; ***, <0.001 and ****, <0.0001.)

| **Data family** | **PC9 PFK-158 plus EGFRi viable cells** | | | | |
|---|---|---|---|---|---|
| **Tukey's multiple comparisons test** | Mean Diff. | 95.00% CI of diff. | Below threshold? | Summary | **Adjusted P Value** |
| DMSO vs. PFK158 | 0.959 | 0.576 to 1.34 | Yes | **** | **<0.0001** |
| DMSO vs. ERLO | 0.913 | 0.666 to 1.16 | Yes | **** | **<0.0001** |
| DMSO vs. ERLO+158 | 1.49 | 1.14 to 1.83 | Yes | **** | **<0.0001** |
| PFK158 vs. ERLO | -0.0456 | -0.338 to 0.247 | No | ns | **0.9644** |
| PFK158 vs. ERLO+158 | 0.527 | 0.156 to 0.897 | Yes | ** | **0.006** |
| ERLO vs. ERLO+158 | 0.572 | 0.304 to 0.841 | Yes | *** | **0.0003** |
| **Tukey's multiple comparisons test** | Mean Diff. | 95.00% CI of diff. | Below threshold? | Summary | **Adjusted P Value** |
| DMSO vs. PFK158 | 0.959 | 0.5764 to 1.342 | Yes | **** | **<0.0001** |
| DMSO vs. OSI | 0.8794 | 0.5342 to 1.225 | Yes | **** | **<0.0001** |
| DMSO vs. OSI+158 | 1.395 | 1.088 to 1.702 | Yes | **** | **<0.0001** |
| PFK158 vs. OSI | -0.07951 | -0.4906 to 0.3315 | No | ns | **0.9354** |
| PFK158 vs. OSI+158 | 0.4359 | 0.03888 to 0.8329 | Yes | * | **0.0306** |
| OSI vs. OSI+158 | 0.5154 | 0.2405 to 0.7903 | Yes | *** | **0.0007** |

| **Data family** | **PC9 KAN757 plus EGFRi viable cells** | | | | |
|---|---|---|---|---|---|
| **Tukey's multiple comparisons test** | Mean Diff. | 95.00% CI of diff. | Below threshold? | Summary | **Adjusted P Value** |
| DMSO vs. KAN757 | 1.5 | 1.20 to 1.80 | Yes | **** | **<0.0001** |
| DMSO vs. ERLO | 0.913 | 0.666 to 1.16 | Yes | **** | **<0.0001** |
| DMSO vs. ERLO+KAN | 1.76 | 1.50 to 2.02 | Yes | **** | **<0.0001** |
| KAN757 vs. ERLO | -0.587 | -0.857 to -0.316 | Yes | *** | **0.0002** |
| KAN757 vs. ERLO+KAN | 0.263 | 0.130 to 0.396 | Yes | *** | **0.0005** |
| ERLO vs. ERLO+KAN | 0.849 | 0.678 to 1.02 | Yes | **** | **<0.0001** |
| **Tukey's multiple comparisons test** | Mean Diff. | 95.00% CI of diff. | Below threshold? | Summary | **Adjusted P Value** |
| DMSO vs. KAN757 | 1.5 | 1.199 to 1.801 | Yes | **** | **<0.0001** |
| DMSO vs. OSI | 0.8794 | 0.5342 to 1.225 | Yes | **** | **<0.0001** |
| DMSO vs. OSI+KAN | 1.798 | 1.509 to 2.086 | Yes | **** | **<0.0001** |
| KAN757 vs. OSI | -0.6205 | -0.9349 to -0.3061 | Yes | *** | **0.0005** |
| KAN757 vs. OSI+KAN | 0.2979 | 0.1246 to 0.4712 | Yes | ** | **0.0015** |
| OSI vs. OSI+KAN | 0.9183 | 0.6366 to 1.200 | Yes | **** | **<0.0001** |

**Table 1S.** Complete statistical analysis data presented for cell viability experiments performed on PC9 cells and presented in Fig.3 D.

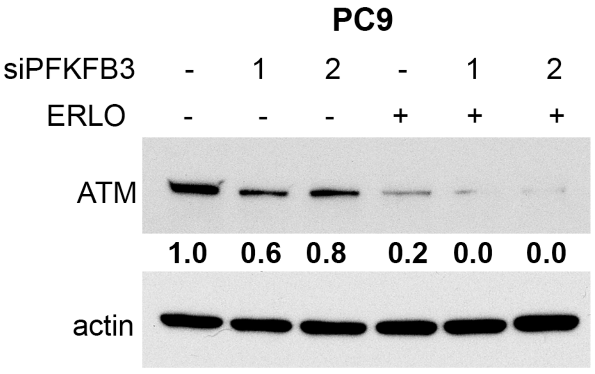

**Fig.2S.** PC9 cells were transfected with PFKFB3 siRNAs (si#1, si#2) followed by treatment with either vehicle or erlotinib (PC9, 0.5 μM) for 24 h. ATM protein levels were analyzed in whole cell lysates by Western blotting. β-actin was used as a loading control. Target ratios were quantified using densitometry and normalized to vehicle-treated samples.

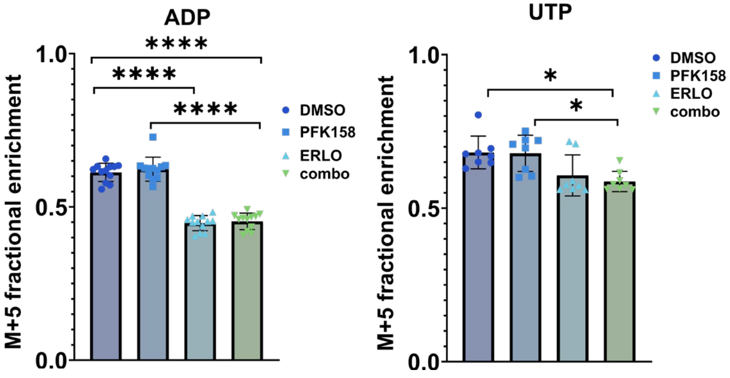

**Fig.3S.** PC9 cells were exposed to appropriate treatments for 36 hours (including 24h exposure to tracer - [U-13C]-glucose.) Fractional enrichment of m + 5 labeled nucleotides and nucleosides presented. Values represent biological replicates from 2 independent experiments (n=8). Data analyzed by TWO-WAY ANOVA with Tukey's *post hoc* tests (p values are shown as follows: *, <0.05; ****, <0.0001.) **B**, Complete statistical analysis data presented for cell viability experiments performed on PC9 cells and presented in Fig.6 C-D (next slide).

| Data family | PC9 PFK-158 plus erlotinib plus NTP viable cells | | | |
|---|---|---|---|---|
| **Tukey's multiple comparisons test** | Mean Diff. | 95.00% CI of diff. | Summary | Adjusted P Value |
| CTRL | | | | |
| DMSO vs. PFK158 | 0.3746 | 0.2282 to 0.5211 | **** | <0.0001 |
| DMSO vs. ERLO | 0.2322 | 0.08576 to 0.3786 | *** | 0.0007 |
| DMSO vs. combo | 0.5061 | 0.3597 to 0.6525 | **** | <0.0001 |
| PFK158 vs. ERLO | -0.1425 | -0.2889 to 0.00395 | ns | 0.0591 |
| PFK158 vs. combo | 0.1315 | -0.01495 to 0.2779 | ns | 0.0921 |
| ERLO vs. combo | 0.2739 | 0.1275 to 0.4204 | **** | <0.0001 |
| NTPs | | | | |
| DMSO vs. PFK158 | 0.401 | 0.2546 to 0.5474 | **** | <0.0001 |
| DMSO vs. ERLO | 0.04583 | -0.1006 to 0.1923 | ns | 0.8356 |
| DMSO vs. combo | 0.3561 | 0.2097 to 0.5025 | **** | <0.0001 |
| PFK158 vs. ERLO | -0.3552 | -0.5016 to -0.2088 | **** | <0.0001 |
| PFK158 vs. combo | -0.04494 | -0.1914 to 0.1015 | ns | 0.8434 |
| ERLO vs. combo | 0.3103 | 0.1638 to 0.4567 | **** | <0.0001 |

| **Šídák's multiple comparisons test** | Mean Diff. | 95.00% CI of diff. | Below threshold? | Summary | Adjusted P Value |
|---|---|---|---|---|---|
| CTRL - NTPs | | | | | |
| DMSO | 0.003111 | -0.1962 to 0.2025 | ns | >0.9999 | |
| PFK158 | 0.02948 | -0.08208 to 0.1410 | ns | 0.9031 | |
| ERLO | -0.1832 | -0.3010 to -0.0655( | ** | 0.0033 | |
| combo | -0.1469 | -0.3897 to 0.09583 | ns | 0.3032 | |

| Data family | HCC827 PFK-158 plus erlotinib plus NTP viable cells | | | | |
|---|---|---|---|---|---|
| **Tukey's multiple comparisons test** | Mean Diff. | 95.00% CI of diff. | Below threshold? | Summary | Adjusted P Value |
| CTRL | | | | | |
| DMSO vs. PFK158 | 0.5607 | 0.4333 to 0.6881 | **** | <0.0001 | |
| DMSO vs. ERLO | 0.5913 | 0.4640 to 0.7187 | **** | <0.0001 | |
| DMSO vs. combo | 0.7797 | 0.6523 to 0.9071 | **** | <0.0001 | |
| PFK158 vs. ERLO | 0.03064 | -0.09675 to 0.1580 | ns | 0.9168 | |
| PFK158 vs. combo | 0.219 | 0.09160 to 0.3464 | *** | 0.0002 | |
| ERLO vs. combo | 0.1884 | 0.06096 to 0.3157 | ** | 0.0016 | |
| NTPs | | | | | |
| DMSO vs. PFK158 | 0.6472 | 0.5198 to 0.7746 | **** | <0.0001 | |
| DMSO vs. ERLO | 0.4524 | 0.3250 to 0.5798 | **** | <0.0001 | |
| DMSO vs. combo | 0.7497 | 0.6223 to 0.8771 | **** | <0.0001 | |
| PFK158 vs. ERLO | -0.1948 | -0.3222 to -0.06740 | ** | 0.0011 | |
| PFK158 vs. combo | 0.1025 | -0.02494 to 0.2298 | ns | 0.1534 | |
| ERLO vs. combo | 0.2972 | 0.1698 to 0.4246 | **** | <0.0001 | |

| **Šídák's multiple comparisons test** | Mean Diff. | 95.00% CI of diff. | Below threshold? | Summary | Adjusted P Value |
|---|---|---|---|---|---|
| CTRL - NTPs | | | | | |
| DMSO | -0.01044 | -0.2098 to 0.1889 | ns | 0.9997 | |
| PFK158 | 0.07606 | -0.05929 to 0.2114 | ns | 0.3538 | |
| ERLO | -0.1494 | -0.2846 to -0.01418 | * | 0.03 | |
| combo | -0.04048 | -0.1578 to 0.07683 | ns | 0.7727 | |

| Data family | PC9 PFK-158 plus erlotinib plus NAC viable cells | | | |
|---|---|---|---|---|
| **Tukey's multiple comparisons test** | Mean Diff. | 95.00% CI of diff. | Summary | Adjusted P Value |
| CTRL | | | | |
| DMSO vs. PFK158 | 0.3746 | 0.2530 to 0.4963 | **** | <0.0001 |
| DMSO vs. ERLO | 0.2322 | 0.1106 to 0.3538 | **** | <0.0001 |
| DMSO vs. combo | 0.5061 | 0.3845 to 0.6277 | **** | <0.0001 |
| PFK158 vs. ERLO | -0.1425 | -0.2641 to -0.02084 | * | 0.0161 |
| PFK158 vs. combo | 0.1315 | 0.009842 to 0.2531 | * | 0.0297 |
| ERLO vs. combo | 0.2739 | 0.1523 to 0.3956 | **** | <0.0001 |
| NAC | | | | |
| DMSO vs. PFK158 | 0.3013 | 0.1797 to 0.4230 | **** | <0.0001 |
| DMSO vs. ERLO | 0.02363 | -0.09800 to 0.1453 | ns | 0.9536 |
| DMSO vs. combo | 0.2782 | 0.1566 to 0.3998 | **** | <0.0001 |
| PFK158 vs. ERLO | -0.2777 | -0.3993 to -0.1561 | **** | <0.0001 |
| PFK158 vs. combo | -0.02315 | -0.1448 to 0.09848 | ns | 0.9562 |
| ERLO vs. combo | 0.2546 | 0.1329 to 0.3762 | **** | <0.0001 |

| **Šídák's multiple comparisons test** | Mean Diff. | 95.00% CI of diff. | Summary | Adjusted P Value |
|---|---|---|---|---|
| CTRL - NAC | | | | |
| DMSO | 2.83E-05 | -0.1402 to 0.1403 | ns | >0.9999 |
| PFK158 | -0.07328 | -0.2432 to 0.09666 | ns | 0.603 |
| ERLO | -0.2085 | -0.3263 to -0.09073 | ** | 0.0013 |
| combo | -0.2279 | -0.3605 to -0.09529 | ** | 0.0017 |

| Data family | HCC827 PFK-158 plus erlotinib plus NAC viable cells | | | |
|---|---|---|---|---|
| **Tukey's multiple comparisons test** | Mean Diff. | 95.00% CI of diff. | Summary | Adjusted P Value |
| CTRL | | | | |
| DMSO vs. PFK158 | 0.5607 | 0.3932 to 0.7282 | **** | <0.0001 |
| DMSO vs. ERLO | 0.5913 | 0.4238 to 0.7589 | **** | <0.0001 |
| DMSO vs. combo | 0.7797 | 0.6122 to 0.9472 | **** | <0.0001 |
| PFK158 vs. ERLO | 0.03064 | -0.1369 to 0.1982 | ns | 0.9608 |
| PFK158 vs. combo | 0.219 | 0.05146 to 0.3865 | ** | 0.0061 |
| ERLO vs. combo | 0.1884 | 0.02082 to 0.3559 | * | 0.0223 |
| NAC | | | | |
| DMSO vs. PFK158 | 0.2914 | 0.1239 to 0.4589 | *** | 0.0002 |
| DMSO vs. ERLO | 0.3181 | 0.1505 to 0.4856 | **** | <0.0001 |
| DMSO vs. combo | 0.3463 | 0.1788 to 0.5138 | **** | <0.0001 |
| PFK158 vs. ERLO | 0.02665 | -0.1409 to 0.1942 | ns | 0.9736 |
| PFK158 vs. combo | 0.0549 | -0.1126 to 0.2224 | ns | 0.816 |
| ERLO vs. combo | 0.02824 | -0.1393 to 0.1958 | ns | 0.9689 |

| **Šídák's multiple comparisons test** | Mean Diff. | 95.00% CI of diff. | Summary | Adjusted P Value |
|---|---|---|---|---|
| CTRL - NAC | | | | |
| DMSO | -0.03465 | -0.2345 to 0.1652 | ns | 0.9749 |
| PFK158 | -0.3039 | -0.4394 to -0.1684 | *** | 0.0007 |
| ERLO | -0.3079 | -0.5025 to -0.1133 | ** | 0.0034 |
| combo | -0.468 | -0.7442 to -0.1919 | ** | 0.0043 |

**Table 2S.** Complete statistical analysis data presented for cell viability experiments performed on PC9 cells and presented in Fig.6 C, D.
